# Succinate dehydrogenase/complex II is critical for metabolic and epigenetic regulation of T cell proliferation and inflammation

**DOI:** 10.1101/2021.10.26.465727

**Authors:** Xuyong Chen, Benjamin Sunkel, Meng Wang, Siwen Kang, Tingting Wang, JN Rashida Gnanaprakasam, Lingling Liu, Teresa A. Cassel, David A. Scott, Ana M. Muñoz-Cabello, Jose Lopez-Barneo, Jun Yang, Andrew N. Lane, Gang Xin, Benjamin Stanton, Teresa W.-M. Fan, Ruoning Wang

## Abstract

Robust and effective T cell-mediated immune responses require the proper allocation of metabolic resources to sustain energetically costly processes like growth, proliferation, and cytokine production. Epigenetic control of the genome also governs T cell transcriptome and T cell lineage commitment and maintenance. Cellular metabolic programs interact with epigenetic regulation by providing substrates for covalent modifications of chromatin. By employing complementary genetic, epigenetic, and metabolic approaches, we revealed that tricarboxylic acid (TCA) cycle flux fuels biosynthetic processes while controlling the ratio of α-ketoglutarate/succinate to modulate the activities of dioxygenases that are critical for driving T cell inflammation. In contrast to cancer cells, where succinate dehydrogenase (SDH)/complex II inactivation drives cell transformation and growth, SDH/complex II deficiency in T cells causes proliferation and survival defects when the TCA cycle is truncated, blocking carbon flux to support nucleosides biosynthesis. Accordingly, replenishing the intracellular nucleoside pool partially relieved the dependence of T cells on SDH/complex II for proliferation and survival. Conversely, SDH deficiency induces a pro-inflammatory gene signature in T cells and promotes T helper 1 (T_H_1) and T helper 17 (T_H_17) lineage differentiation. Mechanistically, the hypoxia-inducible factor 1 (HIF-1) is not required for succinate-induced inflammation in T cells. A reduced α-ketoglutarate/succinate ratio in SDH deficient T cells promotes inflammation through changing the pattern of the transcriptional and chromatin-accessibility signatures and consequentially increasing the expression of the transcription factor, B lymphocyte-induced maturation protein-1 (Blimp-1). Collectively, our studies revealed a critical role of SDH/complex II in allocating carbon resources for anabolic processes and epigenetic regulation in T cell proliferation and inflammation.

## Introduction

T cell activation requires reprogramming of central carbon metabolism that transforms carbon and produces chemical energy through glycolysis, the pentose phosphate pathway (PPP), and the tricarboxylic acid (TCA) cycle to prepare T cells for growth, differentiation, and immune defense (Gerriets and Rathmell, 2012; Pearce and Pearce, 2013; Sena et al., 2013). This conversion is reminiscent of the characteristic metabolic switch during the transformation of normal to cancer cells (Dang, 2013; Pavlova and Thompson, 2016). In parallel, T cell activation is accompanied by a dynamic remodeling of chromatin accessibility and DNA modifications, which determine the transcriptional status of lineage-specifying genes. Recent comparative studies have revealed epigenetic patterns in accessible, poised, and silenced gene loci in different CD4 subsets (Calderon et al., 2019; Gate et al., 2018; Ivanov et al., 2009). Mapped permissive and repressive epigenetic modifications in lineage-restricting transcription factors and lineage-specific cytokine genes indicate that such a bivalent epigenetic state is critical for T cell lineage commitment (Ivanov et al., 2009; Oestreich and Weinmann, 2012; Wilson et al., 2009). As such, a dynamic epigenetic regulation allows progeny cells to maintain lineage-specific transcription programs while retaining some level of plasticity in response to environmental cues. Also, central carbon metabolism allocates carbon inputs to metabolites that are substrates, antagonists, and cofactors of epigenetic-modifying enzymes. Thus, central carbon metabolism is instrumental in establishing and maintaining the epigenetic landscape for cell fate determination. How and which metabolic pathways control the epigenome for the proliferation and differentiation of Teff cells remains elusive.

While an essential function of the TCA cycle is to sustain the oxidative carbon flux to meet energy consumption needs, the TCA cycle also coordinates the carbon input (anaplerosis) and carbon output (cataplerosis) to ensure biosynthetic activities, and as several recent studies suggested, epigenetic regulation (Martinez-Reyes and Chandel, 2020; Murphy and O’Neill, 2018; Owen et al., 2002). Specifically, α-ketoglutarate (α-KG), an intermediate of the TCA cycle, is an essential co-substrate of α-KG dependent dioxygenases such as specific histone and DNA demethylases. In contrast, succinate and fumarate are potent inhibitors of these enzymes (Xiao et al., 2012). Therefore, the ratio changes of these metabolites can impact gene transcription through epigenetic mechanisms (Tsukada et al., 2006; van der Knaap and Verrijzer, 2016; Walport et al., 2012). Succinate dehydrogenase (SDH)/complex II consists of four subunits (SDHA-D) that function in both the TCA cycle and the electron transportation chain (ETC). Its enzymatic activities that convert succinate to fumarate are coupled with reducing FAD to FADH2 and transferring an electron (complex II) into the ETC. Thus, SDH -dependent metabolic reaction represents a critical branch point in the TCA cycle. Intriguingly, SDH is also a tumor-suppressor since inactivating SDH subunits leads to a spectrum of hereditary tumors, suggesting that inactivating this enzyme can confer survival and proliferation advantages to tumor cells (Gottlieb and Tomlinson, 2005; King et al., 2006). As activated T cells share metabolic characteristics with tumor cells (Andrejeva and Rathmell, 2017; Sena et al., 2013), we sought to investigate the role of SDH in T cells. Here, we report that the TCA cycle enzyme SDH is indispensable for driving T cell proliferation by funneling carbon to support anabolic processes. In addition, the TCA cycle flux is constricted based on the availability of succinate that promotes the expression of pro-inflammatory genes through changing the T cell epigenome. Our findings implicate a critical role of the SDH/complex II in coupling the central carbon metabolism to epigenetic regulation to regulate T cell proliferation and inflammation coordinately.

## Results

### SDHB is required for T cell proliferation and survival

Loss-of-function mutations in SDH lead to a spectrum of hereditary tumors, suggesting that the resulting truncated TCA cycle can confer survival and proliferation advantages to tumor cells (Gottlieb and Tomlinson, 2005; King et al., 2006). Active T cells share metabolic characteristics with tumor cells and, therefore, may survive and proliferate in the absence of SDH activities, as a recent study suggested (Bailis et al., 2019). The critical functions of the TCA cycle include producing energy through coupling with the oxidative phosphorylation and generating biosynthetic precursors through cataplerosis(Owen et al., 2002). Accordingly, activated T cells can rapidly incorporate glucose- and glutamine-derived carbons into the TCA cycle metabolites and amino acids, indicating a robust TCA cycle coupled with anaplerosis and cataplerosis (**Fig. S1A-C**). Importantly, SDH inhibitors with distinct modes of inhibition (Ehinger et al., 2016; Miyadera et al., 2003) significantly suppressed proliferation and decreased cell viabilities after T cells activation (**Fig. S2A-C**). To further clarify the role of SDH in T cells, we generated a T cell-specific *SDHB* knockout strain (*SDHB* cKO) by crossing the *SDHB^fl^* strain with the CD4-Cre strain. qPCR and immune blot (IB) analyses validated the deletion of SDHB (**Fig. 1A**), and SDH ablation resulted in a partially truncated TCA cycle, as evidenced by the accumulation of succinate and reduction of fumarate (**Fig. 1B**). While SDHB deletion did not result in any defects in T cell development in the thymus, it did reduce the percentage of T cells, particularly CD8 T cells, in the spleen and lymph node (**Fig. S2D**). In addition, the percentage of naturally occurring interferon- γ(IFN-γ) producing, interleukin-17(IL-17) producing, and Foxp3^+^ CD4 T cells were comparable in both wild-type (WT) and *SDHB* cKO mice (**Fig. S2E**). In contrast to tumor cells, where SDHB inactivation drives cell transformation and growth, SDHB deletion caused more cell death (**Fig.1 E-F, and S2I**) and significantly delayed cell cycle progression from G0/1 to S phase (**Fig.1C**) and suppressed proliferation (**Fig.1 D**) after CD4 T cells activation in vitro, which were associated with moderately reduced cell surface activation markers (**Fig. S2F**), cell size (**Fig. S2G**), and protein content (**Fig. S2H**). To assess the impact of ablating SDHB on CD4 T cells in vivo, we first employed a well-established competitive homeostatic proliferation assay to determine the ratio and carboxyfluorescein succinimidyl ester (CFSE) dilution pattern of purified WT*(Thy1.1^+^)* or *SDHB cKO(Thy1.2^+^)* CD4^+^ T cells in a lymphopenic host (Rag^-/-^). Remarkably, both the ratio between wild type (WT) and *SDHB cKO* CD4^+^ T cells and CFSE dilution patterns suggested that the loss of SDHB dampens T cell proliferation in vivo (**Fig.1G and S2J**). Next, we sought to measure antigen-specific, TCR-dependent proliferation of WT or *SDHB* cKO CD4^+^ T cells. We crossed *Thy1.1* and CD4-Cre, *SDHB^fl^* mice with OT-II transgenic mice to generate WT(*Thy1.1^+^*) and *SDHB cKO(Thy1.2^+^)* donor OT-II strains in CD45.2^+^ background. We then adoptively transferred mixed and CFSE labeled WT and *SDHB* cKO CD4^+^ T cells into CD45.1^+^ mice immunized with chicken ovalbumin (OVA_323-339_). Consistent with the homeostatic proliferation results, *SDHB cKO* OT-II specific CD4^+^ T cells displayed a significant proliferation defect in an antigen-specific manner after immunization (**Fig. 1H and S2K**). Experimental autoimmune encephalomyelitis (EAE), an inflammatory demyelinating disease model, is induced by the proinflammatory T cells in the central nervous system. We used this well-characterized system to investigate the in vivo CD4+ T cell response in *SDHB* cKO mice. In line with our homeostatic and antigen-specific proliferation results, the genetic deletion of SDHB in T cells dramatically reduced leukocyte infiltration and protected the pathogenic progression of mice (**Fig. 1J and 1I**). Collectively, these findings suggest that ablating SDH/complex II activities significantly dampen T cell proliferation and survival in vitro and in vivo.

**Figure 1.**
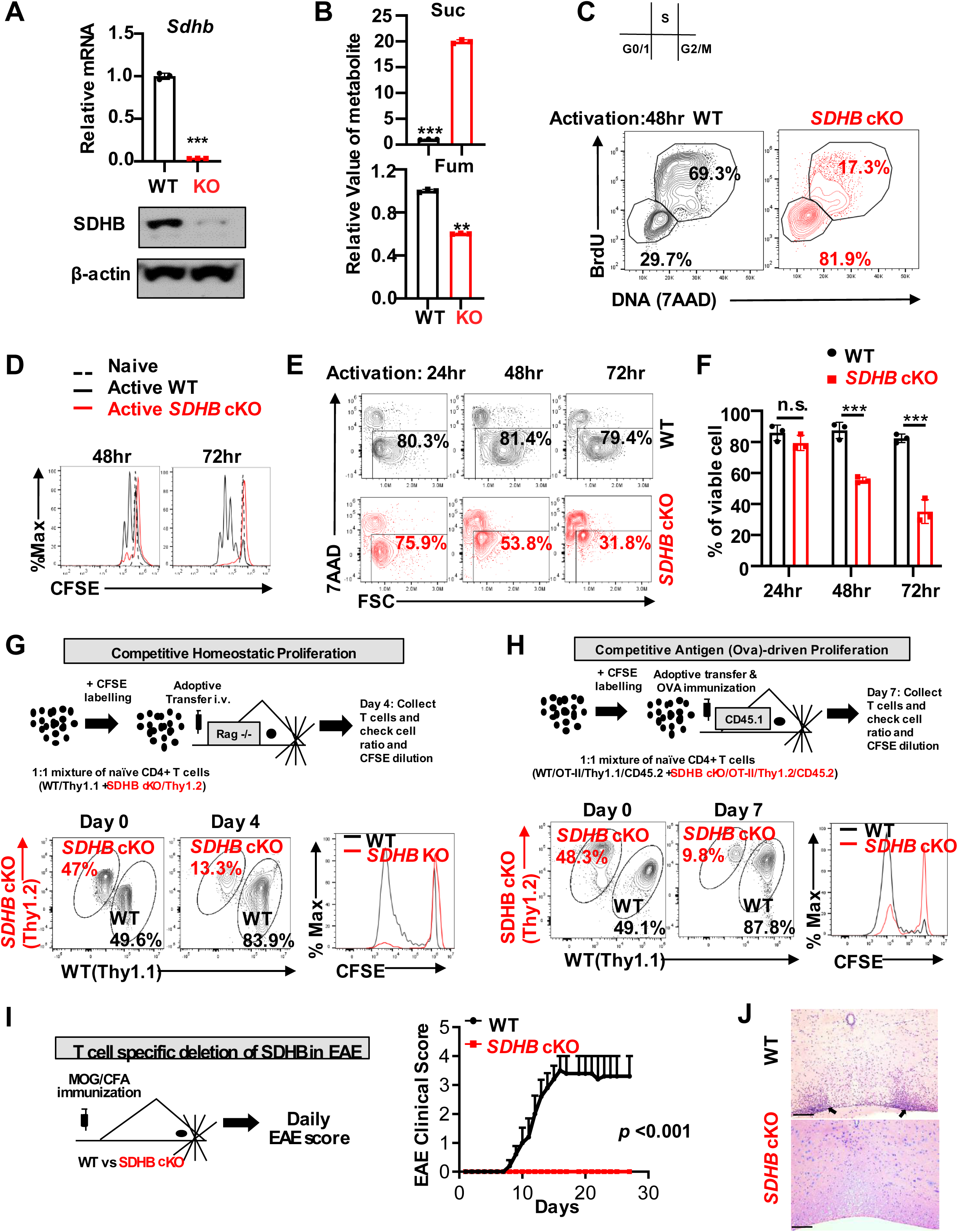
SDHB is required for the T cell proliferation and survival. (**A**) SDHB mRNA and protein levels in activated T cells were determined by qPCR and Immunoblot (mean±SEM, n=3), \*\*\**p* < 0.001, Student’s *t* test. **(B)** Indicated metabolites in activated T cells from WT and *SDHB* cKO mice were determined by CE-QqQ/TOFMS. (mean±SEM, n=3), \*\**p* < 0.01, \*\*\**p* < 0.001, Student’s *t* test. **(C)** The cell cycle of activated CD4^+^ T cells at 48hr was analyzed by using BrdU and 7AAD staining. The numbers indicate the percentage of cells in the cell cycle stage. (**D**) Cell proliferation of naïve and activated CD4^+^ T cells at 48hr and 72hr with indicated genotypes was determined by CFSE dilution. Data represent at least 3 independent experiments. (**E-F**) Cell viability of activated CD4^+^ T cells at 24hr, 48hr, and 72hr with indicated genotypes was assessed by 7AAD staining **(E)**, the statistical analysis of cell viability (mean ± SEM) **(F)**, data represent of 3 independent experiments, \*\*\**p* < 0.001, one-way ANOVA. (**G**) An overview of *in vivo* competitive proliferation experimental procedure (top panel), the donor CD4 T cell ratios before and after adoptive transfer were evaluated by surface staining of isogenic markers, and cell proliferation was evaluated by CFSE dilution (bottle panel), data represent of 3 independent experiments. (**H**) An overview of *in vivo* antigen-specific competitive proliferation experimental procedure (top panel), the donor CD4^+^ T cell ratios before and after adoptive transfer were evaluated by surface staining of isogenic markers, and cell proliferation was evaluated by CFSE dilution (bottle panel), data represent of 3 independent experiments. (**I**) An overview of EAE experimental procedure (left panel), clinical scores were evaluated daily (right panel), data represent mean±SEM (n=5) for each group, experiments were repeated 3 times. **(J)** Hematoxylin and eosin staining of spinal cord sections from WT and *SDHB* cKO mice after EAE induction, leukocyte infiltration is marked by arrowheads. Scale bar, 200 μm.

### SDH inactivation decouples nucleoside biosynthesis from cataplerotic TCA cycle carbon flux

While an essential function of the TCA cycle is to sustain the oxidative carbon flux to meet energy consumption needs, the metabolic flux via the TCA cycle is also critical for carbon allocation, ensuring that biosynthetic precursors production is coupled with nutrient catabolism. Activated *SDHB* cKO T cells consistently displayed lower catabolic activities, such as glycolysis, pentose phosphate pathway (PPP), glutamine oxidation, but not fatty acid oxidation (FAO) as compared with WT active T cells (**Fig. 2A-2D**). Since glutamine is a major anaplerotic substrate of the TCA cycle in T cells after activation (Wang et al., 2011), we supplied ^13^C_5_-glutamine as a metabolic tracer in T cell culture media and followed ^13^C incorporation into downstream metabolites in T cells after activation. The ^13^C_4_ isotopologue of succinate was accumulated, but the corresponding ^13^C_4_ or ^3^C_3_ isotopologues of downstream metabolites including fumarate, aspartate, orotate, and pyrimidine nucleotides were significantly reduced in activated *SDHB* cKO T cells as compared with activated WT T cells (**Fig. 2E**). Similarly, the incorporation of carbon from ^14^C-glutamine into RNA and DNA was significantly suppressed in activated *SDHB* cKO T cells as compared with activated WT T cells (**Fig. 2F**). Metabolite profiling also revealed a global decrease of intracellular pyrimidine nucleotides, purine nucleotides, and their precursors in activated *SDHB* cKO T cells (**Fig. 2G**). Finally, the overall DNA/RNA content was lower in *SDHB* cKO T cells than WT T cells after activation (**Fig. 2H**). Collectively, these results indicate that SDHB deficiency dampens carbon allocation toward nucleotide biosynthesis.

**Figure 2.**
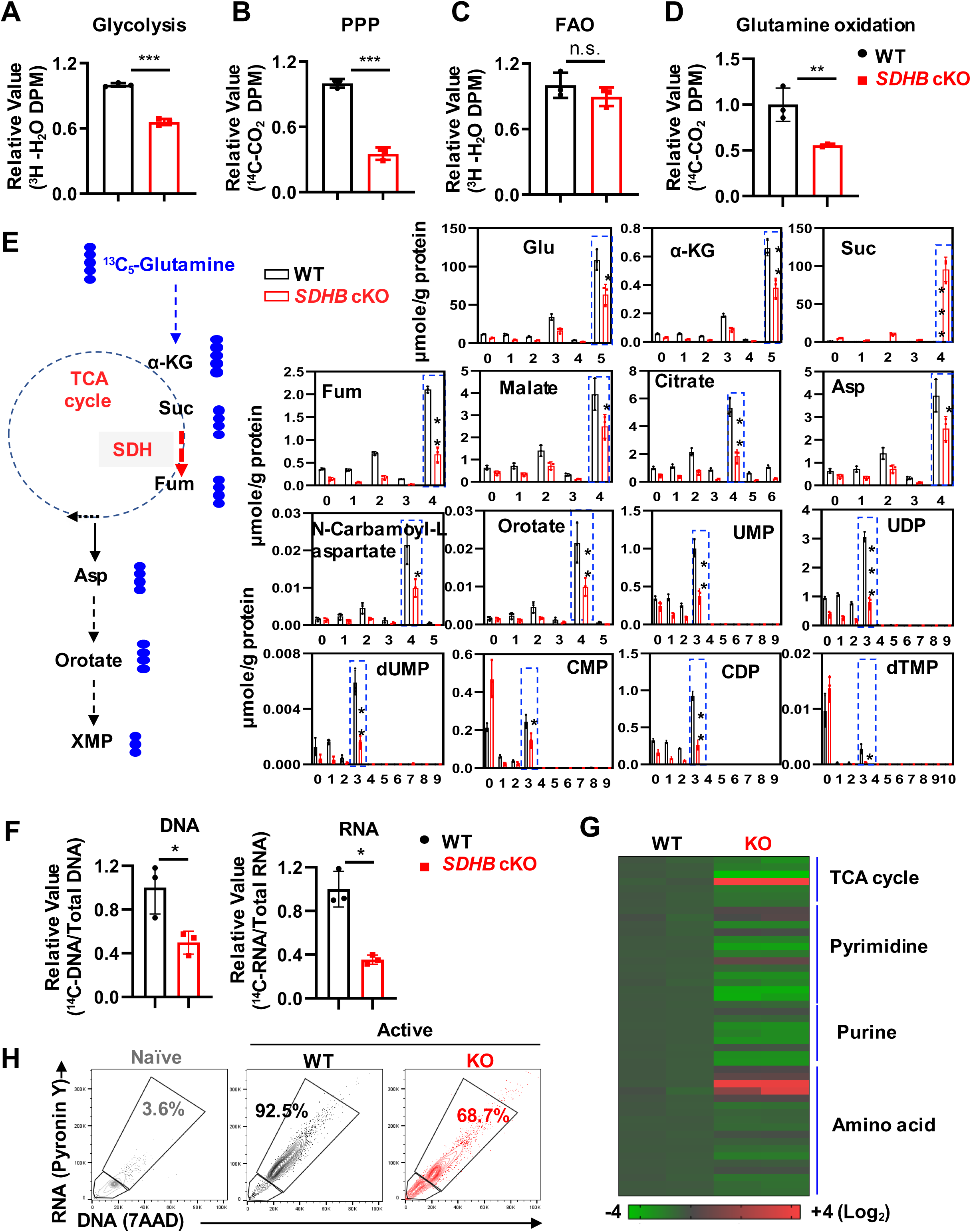
SDHB deficiency decouples the TCA cycle from nucleoside biosynthesis. **(A-D)** Naive CD4^+^ T cells from indicated genotypes were activated for 24 hr, catabolic activities were measured by the generation of ^3^H_2_O from Glucose, D-[5-^3^H(N)] (**A**), ^14^CO_2_ from D-Glucose, [1-^14^C] **(B)**, ^3^H_2_O from Palmitic acid, [9,10-^3^H] (**C**), ^14^CO_2_ from [U-^14^C]-glutamine **(D),** \*\**p* < 0.01, \*\*\**p* < 0.001, Student’s *t* test**. (E)** Diagram of putative catabolic routes of ^13^C_5_-Glutamine in T cells (left panel), metabolites were extracted and analyzed using CE-TOFMS (right panel), numbers in the X-axis represent those of ^13^C atoms in given metabolites, numbers in the Y-axis represent the levels of the metabolites (μmole/g protein). Data represent mean±SEM of (n=3) for each group. **(F)** The incorporation of carbon from ^14^C-glutamine into DNA and RNA was determined by the isotope uptake, \**p* < 0.05, student’s *t* test. **(G)** Indicated metabolites in activated T cells from WT and *SDHB* cKO mice were determined by CE-QqQ/TOFMS. **(H)** DNA and RNA content of naïve and 24 hr active T cells from indicated genotypes were determined by the 7AAD and pyronin-Y uptake.

### Nucleosides supplementation partially compensate for the loss of de novo biosynthesis of nucleotides in *SDHB* cKO T cells

We then sought to determine the importance of SDH in directing TCA cycle carbon flow to nucleotide biosynthesis for supporting T cell proliferation. We reasoned that providing nucleosides can bypass the block in de novo synthesis and consequently overcome some of the defects caused by SDHB deficiency. Indeed, supplementation with a mixture of nucleosides (adenosine, uridine, guanosine, thymidine, cytidine, inosine) partially restored the viability and proliferation of *SDHB* cKO T cells after activation (**Fig. 3A-C**), which was accompanied by partial restoration of DNA/RNA content(**Fig. 3D**) and promote cell cycle progression from G0/1 to S phase (**Fig. 3E**). In addition, adding inosine together with pyrimidines (thymidine, cytidine, and uridine) could maximize the rescue effect (**Fig. 3F-H**), demonstrating the importance of SDH in supporting both purine and pyrimidine biosynthesis.

**Figure 3.**
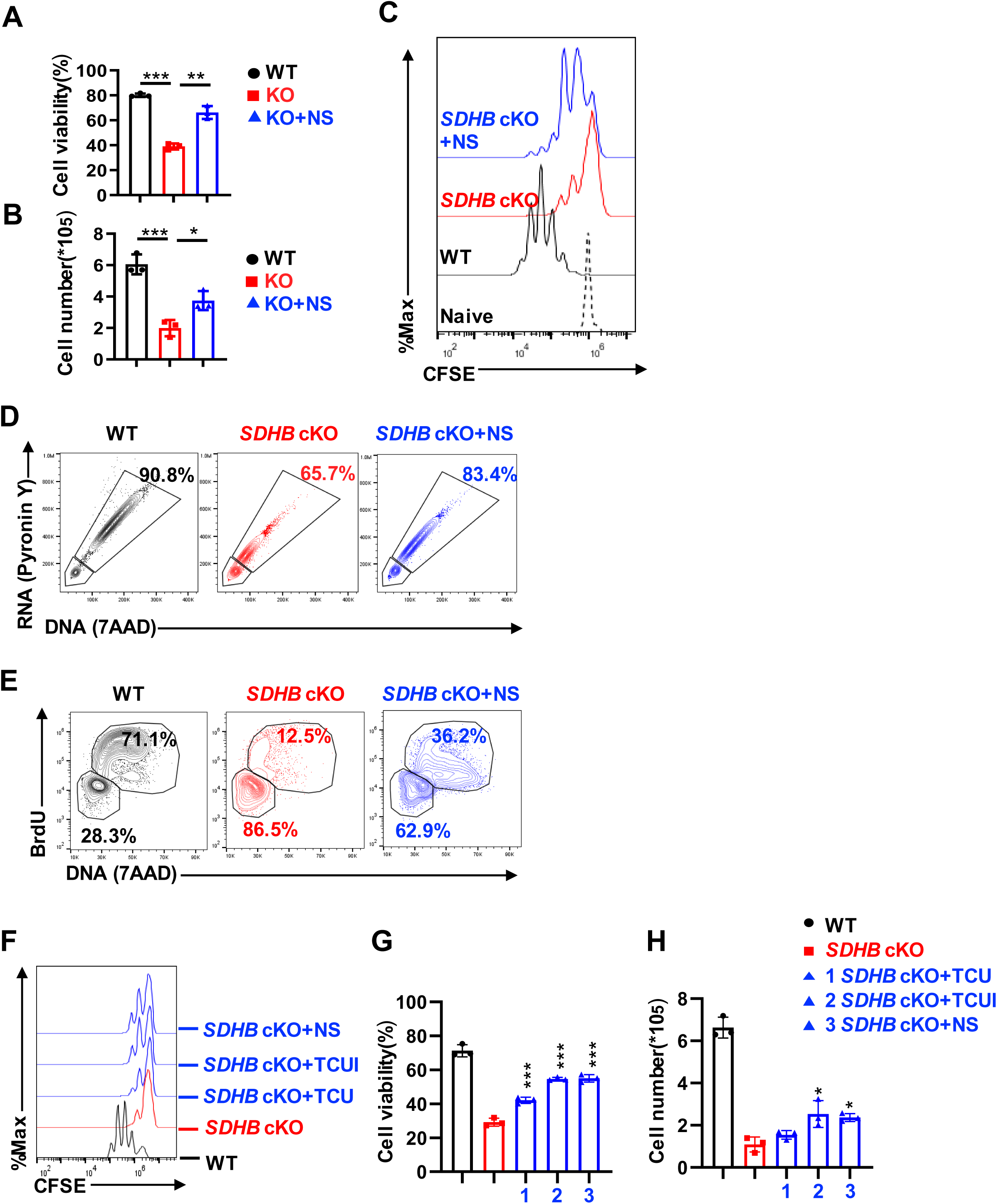
Nucleoside supplementation partially compensate for the loss of de novo biosynthesis of nucleotides in SDHB cKO T cells. **(A-E)** *SDHB* cKO T Cells were supplemented with NS, cell viability at 72 hr was calculated based on the 7AAD staining **(A)**, cell number at 72 hr was measured by a cell counter **(B)**, cell proliferation of naïve and activated CD4^+^ T cells at 72hr was determined by CFSE dilution **(C),** DNA and RNA content were determined by the 7AAD and pyronin uptake **(D)**, the cell cycle of activated CD4^+^ T cells at 48hr was analyzed by using BrdU and 7AAD staining **(E)**. **(F-H)** *SDHB* cKO CD4^+^ T cells were supplemental with mixture nucleosides(NS), inosine together with pyrimidines (TCUI), or pyrimidines(TCU), cell proliferation of activated CD4^+^ T cells at 72hr was determined by CFSE dilution **(F)**, cell viability at 72hr was calculated based on the 7AAD staining **(G)**, cell number at 72 hr was measured by a cell counter **(H)**.Data represent of at least 3 independent experiments, *\*p* < 0.05, *\*\*p* < 0.01, *\*\*\*p* < 0.001, one-way ANOVA.

It is known that nucleotide pool imbalance may cause DNA damage, and consequently, induce apoptosis (Mathews, 2015). In support of this concept, SDHB deletion led to high levels of DNA damage marker (phosphor-histone H2AX), and nucleosides supplementation reversed this effect (**Fig. S3A-B**). Accordingly, the pan-caspase inhibitor (QVD) enhanced the cell viability but not the proliferation of *SDHB* cKO T cells. The combination of nucleosides supplementation and QVD further enhanced the viability but not the proliferation of *SDHB* cKO T cells (**Fig. S3C-D).** Previous studies also suggested that defective SDH/complex II may increase ROS production (Quinlan et al., 2012). However, we only observed a moderate induction of mitochondrial ROS in *SDHB* cKO T cells (**Fig. S3E**). Importantly, ROS scavengers failed to rescue cell viability and proliferation defects in *SDHB* cKO T cells (**Fig. S3F-G**).

To ensure no phenotypic discrepancies between genetic modulation of different subunits of SDH in T cells, we generated a T cell-specific SDHD knockout strain (*SDHD* cKO) by crossing CD4-Cre mice with *SDHD^fl^* mice (Diaz-Castro et al., 2012). Similar to the effect of SDHB deletion in T cells, SDHD deletion reduced the percentage of T cells in the spleen and lymph node without causing defects in T cell development in the thymus (**Fig. S4A-B**). Importantly, SDHD deletion significantly suppressed proliferation and caused more death after T cells activation, which could be partially reversed by nucleosides supplementation (**Fig. S4C-E**). Together, our data suggest that a crucial role of the TCA cycle in supporting T cell proliferation is to allocate carbon for nucleotides biosynthesis (**Fig. S4F**).

### SDH inactivation results in a pro-inflammatory gene signature in T cells after activation

To gain more mechanistic insights into the effects of SDHB deletion on T cells, we performed RNAseq in activated WT and *SDHB* cKO T cells, which revealed an enriched pro-inflammatory gene signature in *SDHB* cKO T cells after activation (**Fig. 4A-C**). Importantly, active but not naïve *SDHB* cKO T cells expressed higher levels of four representative pro-inflammatory genes including *Il17a*, *Il17f*, *Ifng*, *Il22* than WT T cells, indicating that TCR activation is required to achieve an enriched pro-inflammatory gene signature in *SDHB* cKO T cells (**Fig. 4D**). Consistent with gene expression data, the culture media collected from *SDHB* cKO T cells contained significantly higher levels of pro-inflammatory cytokines than the culture media collected from WT T cells (**Fig. 4E**). Nucleosides supplementation, which could partially rescue cell proliferation and survival, failed to reduce the pro-inflammatory cytokine expression in *SDHB* cKO T cells (**Fig. 4F**). Also, the culture media collected from *SDHB* cKO T cells could not increase pro-inflammatory gene expression in WT T cells (**Fig. 4G**). Thus, a cell-intrinsic mechanism that is unrelated to changes in proliferation and survival or secretory molecules is responsible for the pro-inflammatory gene signature in *SDHB* cKO T cells after activation.

**Figure 4.**
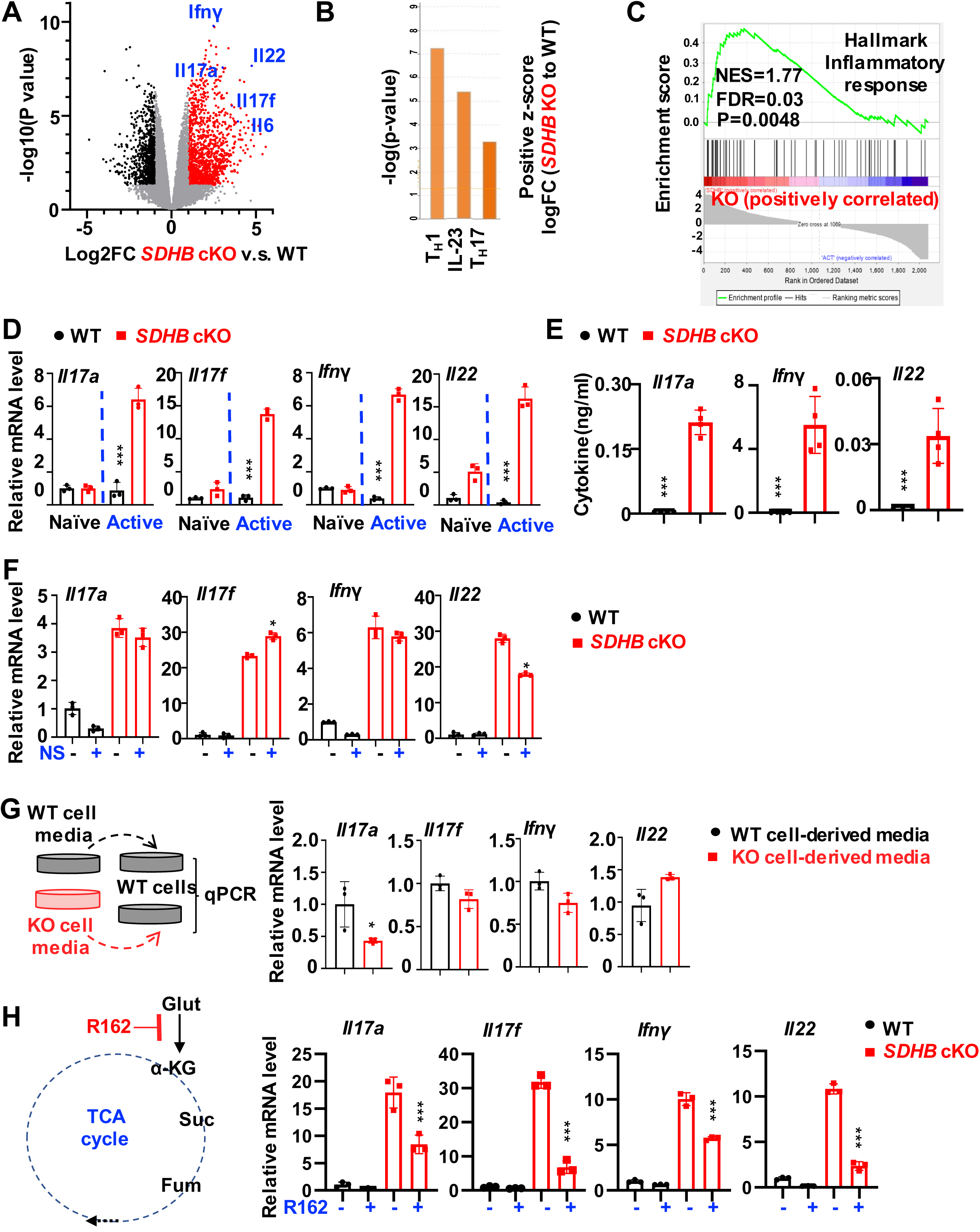
SDHB deficiency promotes a pro-inflammatory gene signature in T cells after activation. **(A)** RAN seq volcano plots of genes differentially expressed in activated CD3 T cells between indicated mouse genotypes (P = 0.05, FC = 2), red color represents enhanced genes, black color represents decreased genes, the blue color indicates presentative genes in the hallmark gene sets. **(B)** Canonical pathway analysis revealed the most upregulated proinflammatory pathways in activated WT and *SDHB* cKO T CD3 T cells cells using ingenuity pathway analysis (IPA)**. (C)** Differentially expressed genes that belong to the enriched hallmark gene sets identified by GSEA are shown in colors, as indicated. **(D)** mRNA levels of *Il17a*, *Il17f*, *Ifng*, *Il22* in naïve and 36hr activated CD4 T cells between indicated mouse genotypes were determined by qPCR (mean±SEM, n=3), \*\*\**p* < 0.001, one-way ANOVA. **(E)** Indicated cytokines in CD4 T cell culture mediums were measured by the LEGENDplex™ kits through flow cytometry, \*\*\**p* < 0.001, student’s *t* test. **(F)** CD4 T cells were activated for 36hr supplemental with or without NS, mRNA levels of indicated genes were measured by qPCR, \**p* < 0.05, one-way ANOVA. **(G)** Experiment procedure of collecting 36hr WT and *SDHB* cKO T cells’ culture medium, then 1:1 ratio mixture with fresh medium and culture WT T cells for 36hr (left panel), cells were collected, and mRNA levels were measured by the qPCR (right panel). \**p* < 0.05, student’s *t* test. **(H)** The diagram of R162 reaction on glutamine-derived anaplerotic flux (left panel), CD4 T cells from indicated genotypes were activated for 36hr with or without R162, mRNA levels were measured by the qPCR (right panel), \*\*\**p* < 0.001, one-way ANOVA.

### Reduced intracellular α-ketoglutarate/succinate ratio increases the expression of pro-inflammatory genes in *SDHB* cKO T cells after activation

SDHB deficiency reciprocally increased the level of succinate while reducing α-KG in T cells after activation (**Fig. 5B**). Such a reduced α-ketoglutarate/succinate ratio can suppress the enzymatic activities of the dioxygenase family, impacting the hypoxia signaling response and DNA/histone methylation pattern (**Fig. 5A**), both of which may lead to a pro-inflammatory response (Eltzschig and Carmeliet, 2011; Palazon et al., 2014; Raghuraman et al., 2016). Correspondingly, cell-permeable succinate partially mimicked the effect of SDHB-deficiency on upregulating pro-inflammatory genes in WT T cells after activation (**Fig. 5C**). Conversely, cell-permeable α-KG reduced the level of pro-inflammatory genes in *SDHB* cKO T cells after activation (**Fig. 5D**). Similarly, a glutamate dehydrogenase inhibitor (R162), which could reduce glutamine-derived anaplerotic flux to attenuate succinate buildup, decreased the expression of pro-inflammatory genes in *SDHB* cKO T cells after activation (**Fig. 4H**). Finally, SDHD deficiency phenocopied SDHB deficiency in inducing pro-inflammatory gene expression, which was reversed by cell-permeable α-KG (**Fig. S4G**).

**Figure 5.**
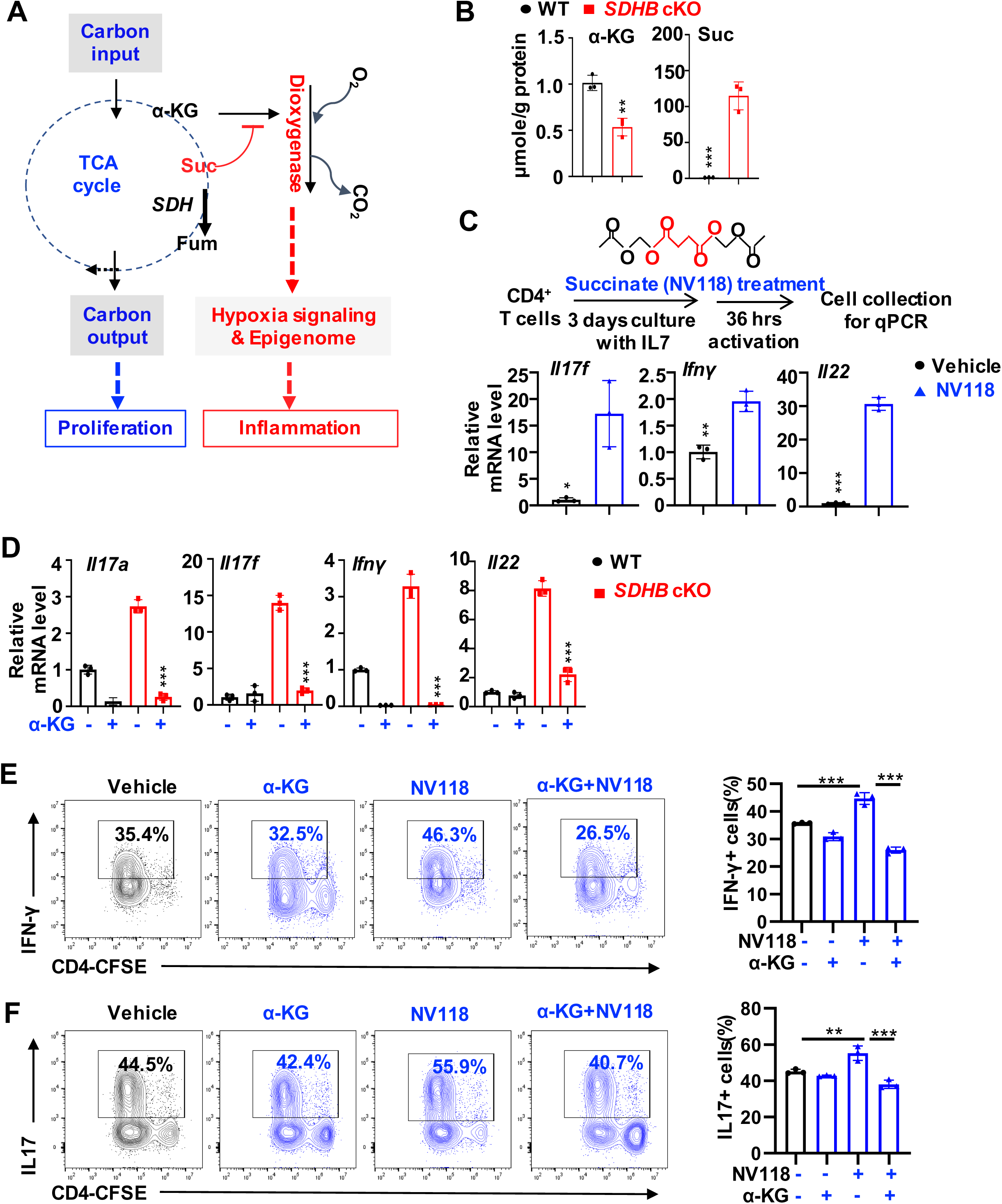
Reduced intracellular α-ketoglutarate/succinate ratio promotes pro-inflammatory signature in T cells after activation. **(A)** Diagram of the putative mechanism of SDHB deficiency induced pro-inflammatory gene signature. **(B)** Indicated metabolites in activated T cells were extracted and analyzed using CE-TOFMS, \*\**p* < 0.01, \*\*\**p* < 0.001, student’s *t* test**. (C)** Structure of NV118, red area represents succinate structure (top panel), the experiment procedure of CD4 T cells culture supplemental with 100μM NV118 (middle panel), mRNA levels of indicated genes were measured by qPCR (bottom panel), \**p* < 0.05, \*\**p* < 0.01, \*\*\**p* < 0.001, student’s *t* test**. (D)** CD4 T cells were activated for 36hr with or without 10mM α-KG, mRNA levels of indicated genes were measured by qPCR (mean±SEM, n=3). **(E-F)** CD4^+^T cells were polarized toward T_H_1 **(E)** and T_H_17 **(F)** lineages with or without 2mM α-KG or 25μM NV118 for 72 hours. The indicated proteins were quantified by intracellular staining. Cell proliferation was determined by CFSE staining. Data represent three independent experiments, \*\**p*<0.01, \*\*\**p*<0.001, one-way ANOVA.

Next, we sought to assess the effects of changing α-ketoglutarate/succinate ratio on CD4 T cell differentiation. Consistent with its effects on enhancing IFN-γ and IL-17 gene expression, cell-permeable succinate enhanced pro-inflammatory T_H_1 and T_H_17 cell differentiation, which could be reversed by cell-permeable α-KG (**Fig. 5E-F**). However, *SDHB* cKO T cells failed to proliferate and could not generate IFN-γ^+^ CD4^+^ T cells under T_H_1 polarization condition or IL-17^+^ CD4^+^ T cells under T_H_17 polarization condition (**Fig. S5A**). The result is in line with previous observations that cell proliferation is required to achieve optimized T cell polarization (Bird et al., 1998). Genomic deletion and RNA depletion of SDHB require a specific time after Cre activation. Also, the protein half-life of SDHB is around 10 hr (Yang et al., 2012). We envisioned that a tamoxifen-inducible Cre recombinase (CreERT2) model might allow us to bypass the effect of SDHB deficiency on T cell activation and proliferation if Cre functions during activation. Compared with WT cells, acute deletion of SDHB largely bypassed its requirement for driving cell proliferation but enhanced T_H_1 and T_H_17 differentiation (**Fig. S5B-C**). Importantly, cell-permeable α-KG could eliminate the effects of acutely deleting SDHB on T cell differentiation (**Fig. S5B-C**). Using this genetic model, we have differentiated the role of SDH in regulating T cell activation and proliferation from its role in driving T cell differentiation. Together, our results suggest that reducing the α-ketoglutarate/succinate ratio promotes pro-inflammatory gene expression and enhances T_H_1 and T_H_17 differentiation in vitro.

### HIF1-α is dispensable for enhanced pro-inflammatory gene signature in *SDHB* cKO T cells

A reduced α-ketoglutarate/succinate ratio suppresses enzymatic activities of the α-KG -dependent dioxygenases, which include prolyl hydroxylases, Ten-eleven translocation (TET) enzymes, and lysine demethylases, and may consequently impact inflammation via HIF-1a and/or epigenetic mechanism (**Fig. 5A**) (Lee et al., 2020; Semenza, 2012). We (and others) have previously shown that HIF1-α plays an essential role in enhancing inflammation in T cells and other immune cells (Dang et al., 2011; Liu et al., 2016; Tannahill et al., 2013). To determine if HIF1-α is responsible for upregulating pro-inflammatory genes, we generated a T cell-specific *SDHB* and *HIF1-α* double knockout (dKO) strain by crossing the *HIF1-α^fl^* with CD4-Cre, *SDHB^fl^* strain. *HIF1-α/SDHB* dKO T cells displayed comparable phenotypes in T cell development, cell survival and proliferation compared with *SDHB* cKO T cells after activation (**Fig. S6A-E**). Importantly, pro-inflammatory gene expression was not reduced in *HIF1-α/SDHB* dKO T cells compared to *SDHB* cKO T cells after activation (**Fig. S6F**). Conversely, cell-permeable succinate could enhance T_H_1 and T_H_17 cell differentiation in a HIF1-α independent manner (**Fig. S6G-H**). Collectively, our results suggest that HIF1-α does not mediate pro-inflammatory phenotypes in *SDHB* cKO T cells after activation.

### SDHB deficiency promotes a pro-inflammatory gene signature through an epigenetic mechanism

Next, we measured succinate, α-ketoglutarate, DNA, and histone methylation levels in WT and *SDHB* cKO naïve T cells. Compared with the WT T cells, the *SDHB* cKO naïve T cells contained a higher level of succinate but not α-KG than WT T cells (**Fig. S7A-B**). In contrast to control T cells, the *SDHB* cKO T cells or T cells treated with cell-permeable succinate displayed higher levels of 5-methylcytosine (5-mC) and histone methylations in several key lysine residues that are often associated with transcriptional activities (Kouzarides, 2007; Wu and Zhang, 2017) (**Fig. S7C-F**). We reasoned that the DNA and histone methylations might work in concert to reprogram an epigenetic state via modulating chromatin accessibility, consequently leading to induction of inflammatory genes in activated T cells with succinate accumulation. To test this hypothesis, we examined the genome-wide transcriptomes and chromatin-accessibility by performing RNAseq and Assay for Transposase Accessible Chromatin with high-throughput sequencing (ATAC-seq) in parallel in activated CD4^+^ WT and *SDHB* cKO T cells. Indeed, ATAC-seq revealed distinctive patterns of DNA accessibility that are associated with genomic loci of genes involved in T cell activation and inflammation in activated WT and *SDHB* cKO CD4 T cells (**Fig. 6A-B**). After integrating the RNA-seq and ATAC-seq data, we found that the induction of pro-inflammatory genes and transcription factors known to regulate T cell differentiation and inflammation were concordant with the increase in chromatin accessibility of these genes in the activated *SDHB* cKO CD4 T cells (**Fig. 6C**). *PRDM1* was among these genes with increased transcript and DNA accessibility. *PRDM1* encodes B-lymphocyte-induced maturation protein (BLIMP1), a transcription factor known which directly bound and co-localized with STAT-3, p300, RORγt on the Il23r, Il17f, and Csf2 cytokine loci to regulate T cell differentiation and inflammation (Gaffen et al., 2014; Jain et al., 2016). Importantly, analysis of transcriptional factor binding motifs in the differential accessibility regions revealed that PRDM1 was one of the top transcription factors that may gain increased chromatin accessibility at the regulatory elements of the T cell activation and inflammation genes in the activated *SDHB* cKO CD4 T cells (**Fig. 6D**). In addition, ATAC-seq analysis of T cells of WT and *SDHB* cKO under naïve conditions also showed a significant motif enrichment of *PRDM1* in the differential accessibility regions, portending the increased transcription of its target gene upon stimulation (**Fig. S7G-H**). We, therefore, hypothesized that transcription factors, such as PRDM1 that showed concordant increases in both their expression level and DNA accessibility, may play a key role in driving inflammation in *SDHB* cKO CD4 T cells. Among a panel of transcription factors that displayed enhanced DNA accessibility in either activated or naïve *SDHB* cKO CD4 T cells (**Fig. 6D and S7H)**, only PRDM1’s expression was enhanced by cell-permeable succinate and SDHB deletion (**Fig. 6E-G)**. Moreover, cell-permeable α-KG reduced the expression of PRDM1 in *SDHB* cKO CD4 T cells (**Fig. 6E-F**). Next, we employed a CRISPR-Cas9 approach to testing if *PRDM1* is required for regulating the expression of inflammatory genes and T cell differentiation in the context of SDHB deletion and succinate accumulation. Indeed, deletion of *PRDM1* reduced the level of pro-inflammatory genes *Il17a*, *Il17f*, *Ifng*, but not *Il22* in *SDHB* cKO T cells (**Fig. 7A-B)**. Moreover, deletion of *PRDM1* eliminated the effect of cell-permeable succinate on promoting T_H_1 and T_H_17 differentiation (**Fig. 7C-D**). Collectively, these results suggest that TCA cycle metabolites (α-KG/succinate) regulate T cell inflammation through a coordinated epigenetic and transcriptional control of inflammatory genes.

**Figure 6.**
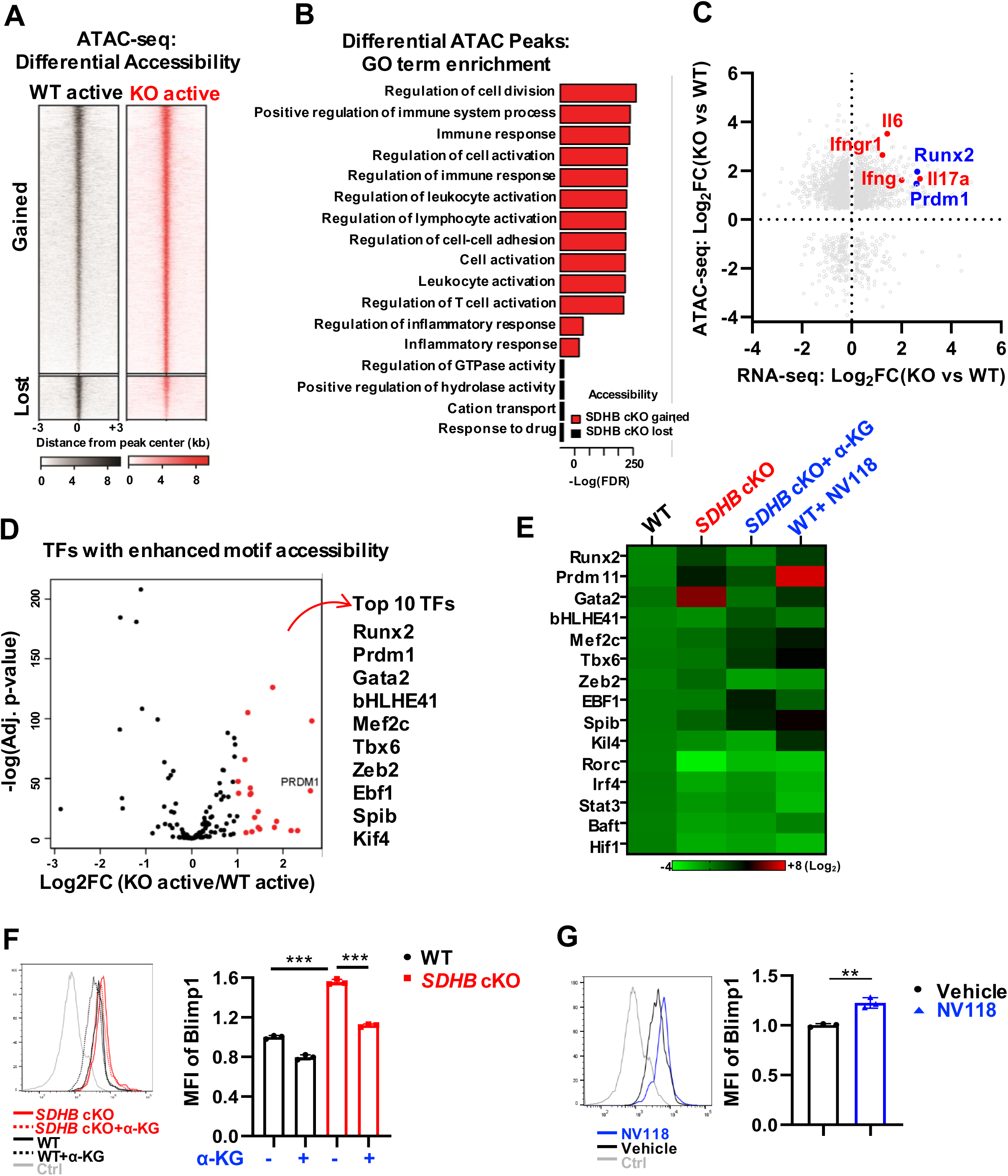
SDHB deficiency increases succinate to enhance DNA accessibility and pro-inflammatory genes transcription. (**A**) Differential chromatin accessibility was measured by ATAC-seq in activated CD4 T cells between indicated genotypes (n = 3 replicates for each genotype), identifying 8,543 sites with accessibility gain and 1,521 sites with accessibility loss in activated *SDHB* cKO T cells. (**B**) ATAC-seq peaks with differential accessibility were linked to nearby genes, and ontology analysis was performed using GREAT. (**C**) Accessibility changes in differential ATAC-seq peaks were plotted against expression changes (CD4 T cells RNA-seq) in the nearby genes identified in (**B**). Concordant changes (i.e., enhanced expression and accessibility) were observed among pro-inflammatory genes and master transcription factors (TFs) mediating inflammation. (**D**) Motif analysis was performed on ATAC-seq peaks showing enhanced accessibility in activated *SDHB* cKO CD4 T cells. A volcano plot shows up-regulated expression of many cognate TFs by RNA-seq. **(E)** Heatmap of selected TFs ware determined in the activated WT and *SDHB* cKO CD4 T cells with indicated treatment. **(F)** Blimp1 protein levels from 36hrs activated WT and *SDHB* cKO CD4 T cells with indicated treatment were measured by intracellular staining through flow cytometry, MFI was analyzed, \*\*\**p*<0.001, one-way ANOVA. **(G)** WT CD4^+^ T cells were incubated without or with NV118 for 72hr in naïve condition then activated for 36hrs without or with NV118. Blimp1 protein levels were measured by intracellular staining through flow cytometry, MFI was analyzed, \*\**p*<0.01, student’s *t* test.

**Figure 7.**
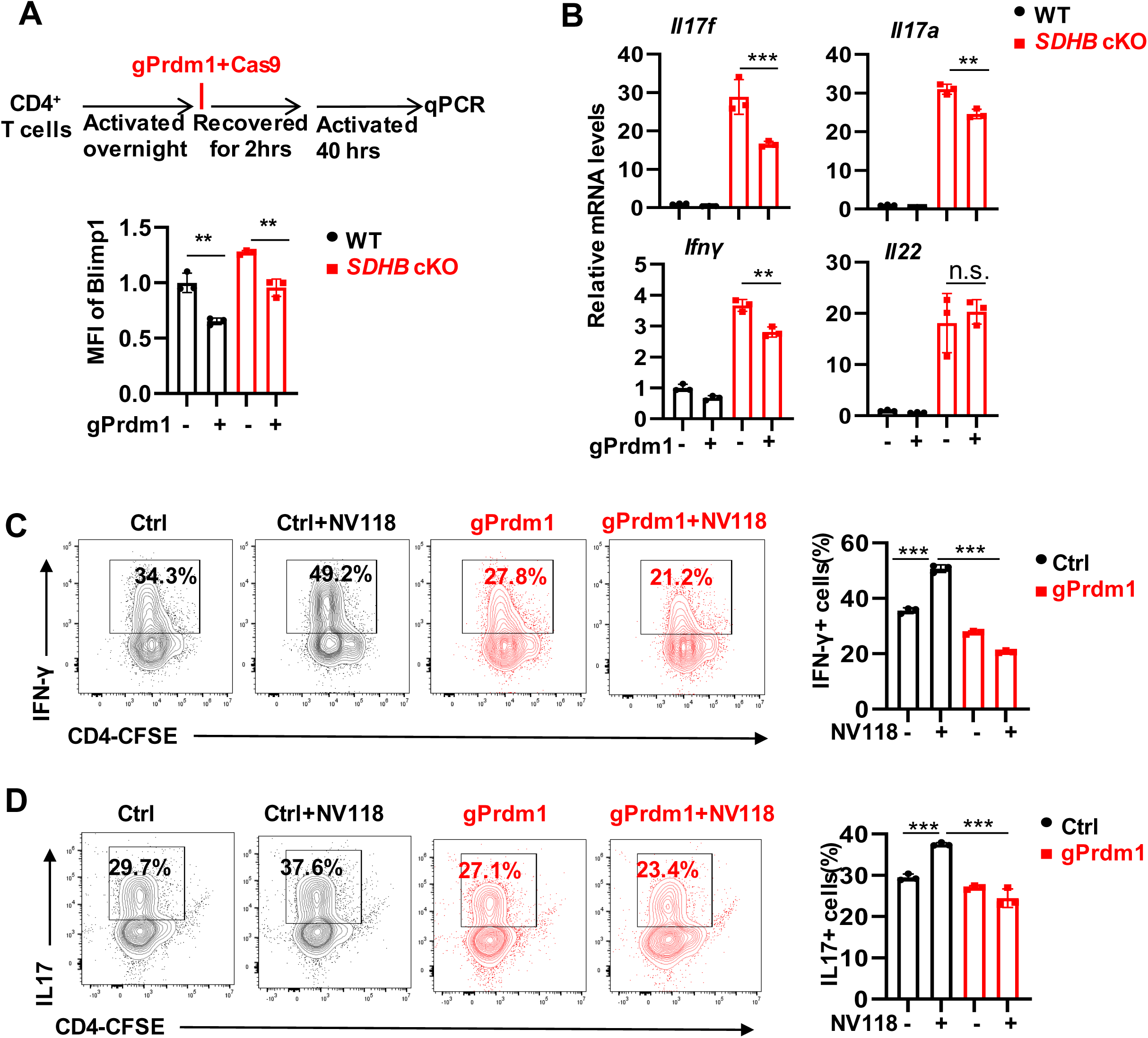
Succinate-mediated Prdm1/Blimp1 expression contributes to proinflammatory signature in T cells. **(A)** Experiment procedure of deletion prdm1 by using CRISPR-CAS9, CD4 T cells were activated overnight, then electroporated with gPrdm1 and Cas9, cells were recovered in culture medium for 2 hrs prior to activation for 40hrs (top panel), Blimp1 protein levels were measured by intracellular staining through flow cytometry (bottle panel), MFI was analyzed, *\*\*p* < 0.01, one-way ANOVA. **(B)** CD4 T cells from indicated genotypes were electroporated with gPrdm1 and Cas9, mRNA levels of indicated genes were determined by qPCR, *\*\*p* < 0.01, *\*\*\*p* < 0.001, one-way ANOVA. **(C-D)** CD4 T cells were activated overnight, then electroporated with gPrdm1 and Cas9. Cells were then polarized toward T_H_1 **(C)** and T_H_17 **(D)** lineages with indicated treatment for 72 hours. The indicated proteins were quantified by intracellular staining, cell proliferation was determined by CFSE staining. Data represent three independent experiments, *\*\*\*p* < 0.001, one-way ANOVA.

## Discussion

Rapidly evolving pathogens impose selective pressures on host immune cells’ metabolic fitness and metabolic plasticity, allowing immune cells to maintain homeostasis while remaining ready to mount rapid responses under diverse metabolic and immune conditions (Slack et al., 2015; Troha and Ayres, 2020; Weinberg, 1975; Weis et al., 2017). Beyond this, the availability of specific metabolites and the pathways that process them interconnect with signaling events and epigenetic regulators in the cell, orchestrating metabolic checkpoints that influence T cell activation, differentiation, and immune function (Birsoy et al., 2015; Chi, 2012; Gerriets and Rathmell, 2012; Michalek and Rathmell, 2010; Pearce and Pearce, 2013; Phan et al., 2017; Powell and Delgoffe, 2010; Russ et al., 2013; Sena et al., 2013). While the essential function of the TCA cycle is to sustain the oxidative carbon flux to meet energy consumption needs, the metabolic flux via the TCA cycle is also critical for carbon anabolism, ensuring that production of biosynthetic precursors of protein, lipids, and DNA/RNA is coupled with energy production from nutrients to sustain cell growth. Accordingly, the major role of the ETC in supporting cell proliferation is to regenerate nicotinamide adenine dinucleotide (NAD)+ and to permit aspartate synthesis in cancer cells (Birsoy et al., 2015; Gaude et al., 2018; Gui et al., 2016; Sullivan et al., 2015). Similarly, we found that carbon input (mainly through glutamine) to replenish TCA cycle intermediates (anaplerosis) is balanced with an output of 4-carbon intermediates from the TCA cycle to aspartate and nucleosides and nucleotides (cataplerosis) in T cells. These results suggest that active T cells share metabolic characteristics with tumor cells (Andrejeva and Rathmell, 2017; Sena et al., 2013). Unlike hyper-proliferative tumor cells where inactivating the TCA cycle enzymes SDH can confer survival and proliferation advantages to tumor cells (Gottlieb and Tomlinson, 2005; King et al., 2006), we found that SDH-dependent steps are vital for carbon allocation via the TCA cycle in supporting anabolic processes to drive T cell proliferation. During transformation, additional genetic alteration may confer a high degree of metabolic adaptation to SDH/complex II deficient cancer cells (Cardaci et al., 2015; Lussey-Lepoutre et al., 2015).

Recent emerging evidence has shown that succinate is necessary to promote inflammation during macrophage activation (Artyomov et al., 2016; Mills and O’Neill, 2014). Endogenous metabolite itaconate regulates inflammatory cytokine production during macrophage activation through targeting SDH/complex II, implicating metabolites as key signals of regulating inflammation (Lampropoulou et al., 2016; Mills et al., 2016; Swain et al., 2020). In macrophages, succinate exerts its pro-inflammatory actions through a plethora of mechanisms, including stabilizing HIF-1α, affecting mitochondrial ROS (mtROS) production, and activating cell surface receptor, the G-protein coupled receptor succinate receptor 1 (SUCNR1) (He et al., 2004; Mills et al., 2016; Ryan et al., 2019; Tannahill et al., 2013). In T cells, the TCA cycle allocates carbon inputs to modulate substrates, antagonists, and cofactors of epigenetic-modifying enzymes, thereby enacting epigenetic changes during T cell activation (Tsukada et al., 2006; van der Knaap and Verrijzer, 2016; Walport et al., 2012). Particularly, succinate is a critical metabolite regulator of histone and DNA demethylases (Martinez-Reyes and Chandel, 2020; Tsukada et al., 2006; van der Knaap and Verrijzer, 2016; Walport et al., 2012). T cell activation is accompanied by a dynamic remodeling of chromatin, accessibility, and histone modifications, which determine the transcriptional signatures of lineage-specifying genes via cell type-specific gene regulatory elements. The epigenetic landscape promotes the faithful execution of lineage-restricting transcription programs while maintaining a degree of plasticity in response to intracellular and environmental cues (Oestreich and Weinmann, 2012; Wilson et al., 2009). Mitochondrial complex III regulates T cell activation and regulatory T cell functions through controlling mitochondria ROS and DNA methylation, respectively (Sena et al., 2013; Weinberg et al., 2019). We uncovered distinctive patterns of DNA accessibility that are associated with genomic loci of inflammatory genes and transcription factors that regulate inflammatory genes in WT and *SDHB* cKO CD4 T cells. By acting on genomic regions where specific subsets of genes locate, transcription factors often coordinate with epigenetic regulators to define cell-and tissue-specific gene expression patterns and consequently determine cell fate. Intriguingly, we observed a concordant increase in the expression and the motif accessibility of transcription factor *PRDM1*, a known regulator of T cell differentiation and inflammation, in *SDHB* cKO CD4 T cells compared to WT CD4 T cell (Gaffen et al., 2014). Deletion of *PRDM1* eliminates the effect of succinate on promoting the expression of the inflammatory genes and effector T cell differentiation. These results support the idea that SDH/complex II is necessary for promoting T cell inflammation through a coordinated epigenetic and transcriptional control of inflammatory genes. Collectively, TCA cycle flux and the ETC may act concertedly to ensure coordinated carbon assimilation, energy production, and epigenetic regulation. The participation of the master transcription factor in such combinational contort adds one layer of regulatory flexibility on cells, permitting contextual gene expression patterns. This mechanism can potentially be generalized to explain how metabolic constraints impose on the epigenetic and transcriptional regulation of a broad range of cell-fate transitions.

## Data availability statement

The CD3 T cells RNA-seq datasets generated for this study can be found in the GEO accession GSE184010. The CD4 T cells RNA-seq datasets generated for this study can be found in the GEO accession GSE184734. The CD4 T cells ATAC-seq datasets generated for this study can be found in the GEO accession GSE184744.

## Author contributions

R. Wang conceptualized and supervised this work. X.C. carried out most of the experiments. B.S., B.S., and M.W. performed and analyzed the CD4 T cells ATAC seq and RNA seq data. S.K., T.W., J.R.G., L.L., T.A.C., D.A.S., Y.Y., A.N.L., and G.X. were involved in data collection, analysis, and review. B.S., and T.W.F., provided conceptual input into the study development. A.M.MC. and J.LB provided some experimental animals. X.C., B.S., A.N.L., T.W.F., and R. Wang wrote the manuscript. All authors discussed the results and provided feedback on the manuscript.

## Materials and Methods

### Mice

Mice with one targeted allele of *SDHB* on the C57BL/6 background (*SDHB*^tm1a(EUCOMM)Hmgu^) were generated by The European Conditional Mouse Mutagenesis Program (EUCOMM) (Skarnes et al., 2011). The mice were first crossed with a transgenic Flippase strain (B6.129S4*Gt(ROSA)26Sor^tm1(FLP1)Dym^*/RainJ) to remove the LacZ-reporter allele and then crossed with the CD4-Cre strain to generate T cell-specific *SDHB* knockout strain (*SDHB* cKO). *SDHB^fl^* were crossed with ROSA26CreERT2 to generate the acute deletion model. *SDHB^fl^* were crossed with *HIF1-α^fl^* (JAX) and CD4-Cre strain to generate *SDHB* and *HIF1-α* double KO (dKO) strain. Mice with one targeted allele of *SDHD* on the C57BL/6 background were crossed with the CD4-Cre strain to generate the CD4Cre *SDHD* cKO mice (*SDHD* cKO) (Diaz-Castro et al., 2012). Mice with a targeted allele of *HIF1-α* on the C57BL/6 background were crossed with the CD4-Cre strain to generate the CD4Cre *HIF1-α* KO mice (*HIF1-α* cKO) (Ryan et al., 2000). OT-II mice (B6.Cg-Tg(TcraTcrb)425Cbn/J) were crossed with CD4Cre *SDHB* cKO mice to generate the OT-II CD4Cre *SDHB* cKO mice. OT-II mice (B6.Cg-Tg(TcraTcrb)425Cbn/J) were crossed with *Thy1.1*^+^ mice (B6.PL-*Thy1*a/CyJ) to generate OT-II *Thy1.1* mice. C57BL/6 (WT) mice, CD45.1^+^ mice (B6.SJL-Ptprca Pepcb/BoyJ), and Rag1-/- mice (B6.129S7-Rag1tm1Mom/J) were obtained from the Jackson Laboratory (JAX, Bar Harbor, ME). Mice with gender and age-matched (7-12 weeks old) were used in the experiments. All mice were bred and kept in specific pathogen-free conditions at the Animal Center of Abigail Wexner Research Institute at Nationwide Children Hospital. Animal protocols were approved by the Institutional Animal Care and Use Committee of the Abigail Wexner Research Institute at Nationwide Children’s Hospital (IACUC; protocol number AR13-00055).

### Mouse T cell isolation and culture

Total CD3^+^ T cells or naïve CD4^+^ T cells were enriched from mouse spleen and lymph nodes by negative selection using MojoSort Mouse CD3 T Cell Isolation Kit or MojoSort Mouse CD4 Naive T Cell Isolation Kit (MojoSort, Biolegend) following the manufacturer’s instructions. Isolated T cells (1×10^6^ cells/mL) were then maintained in culture media containing mouse IL-7 (5 ng/mL) for naïve condition or stimulated with mouse IL-2 (100 U/mL) as well as plate-bound anti-mCD3 (clone 145-2C11) and anti-mCD28 (clone 37.51) antibodies for active condition. Culture plates were pre-coated with 2 μg/mL anti-mCD3 (clone 145-2C11, Bio X Cell) and 2 μg/mL anti-mCD28 (clone 37.51, Bio X Cell) antibodies overnight at 4°C. For T cell’s CFSE dilution analysis, 1-2 x 10^7^ cells were incubated in 4 µM PBS-diluted CFSE (Invitrogen) for 10 min; cells were then washed three times with RPMI-1640 medium containing 10% FBS for culture.

For CD4 T cell differentiation, 48 wells culture plates were pre-coated with 5 μg/mL anti-mCD3 (clone 145-2C11, Bio X Cell) and 5 μg/mL anti-mCD28 (clone 37.51, Bio X Cell) antibodies over two nights at 4°C for T_H_1 and iTreg polarization, and were pre-coated with 10 μg/mL anti-mCD3 (clone 145-2C11, Bio X Cell) and 10 μg/mL anti-mCD28 (clone 37.51, Bio X Cell) antibodies over two nights at 4°C for T_H_17 polarization. Freshly isolated naïve CD4^+^ T cells (0.6 ×10^6^/mL) were stimulated with mIL-2 (200 U/mL), mIL-12 (5 ng/mL, PeproTech) for T_H_1 differentiation, were stimulated with mIL-6 (50 ng/mL, PeproTech), hTGF-β1 (2 ng/mL), anti–mIL-2 (8 μg/mL, Bio X Cell), anti–mIL-4 (8 μg/mL, Bio X Cell), and anti–mIFN-γ (8 μg/mL, Bio X Cell) for T_H_17 differentiation, and were stimulated with mIL-2 (200 U/mL) and hTGF-β1 (5 ng/mL, PeproTech) for iTreg differentiation. At day 3 or day 4, eBioscience™ Cell Stimulation Cocktail of phorbol 12-myristate 13-acetate (PMA), ionomycin, brefieldin A and monensin (eBioscience) were added into T_H_1 or T_H_17 cells for induction and subsequent intracellular detection of cytokines for four hours and then collected the cells for fixation and staining.

All cells were cultured in RPMI 1640 media supplemented with 10% (v/v) heat-inactivated dialyzed fetal bovine serum (DFBS), 2 mM L-glutamine, 0.05 mM 2-mercaptoethanol, 100 units/mL penicillin and 100 µg/mL streptomycin at 37°C in 5% CO2.

A mixture of nucleosides (NS) including Adenosine, Uridine, Inosine, Cytidine, Guanosine, Thymidine (Sigma) each at 50uM or TCUI combination (Thymidine, Cytidine, Uridine, Inosine) or TCU (Thymidine, Cytidine, Uridine), Atpenin A5 (A5, 100nM), diacetoxymethyl malonate (NV161,100μM), dimethyl-itaconate (DI, 0.5mM), or 4-octyl-itaconate (4OI, 500μM), 20uM QVD-OPh, 1mM NAC, 1mM GSH EE, 100μM or 25 μM NV118, 10mM or 2mM dimethyl α-KG, 50μM R162, 600nM 4OHT were used in the indicated experiments, cell culture related cytokines, antibodies, inhibitors and chemicals were list in table 1.

### Flow Cytometry

For analysis of surface markers, cells were suspended in phosphate-buffered saline (PBS) containing 2% (w/v) bovine serum albumin (BSA) and incubated with antibodies (Table 2) for 30 min at 4°C. Cells were washed and stained with 7AAD (Biolegend) for cell viability analysis for 5 min before running on a flow cytometer.

For intracellular cytokine staining, after Cell Stimulation Cocktails incubation, cells were collected and stained with surface markers, then fixed and permeabilized using FoxP3 Fixation/Permeabilization Kit (eBioscience), cells were stained with anti-IFN-g or anti-IL-17A antibodies (Biolegend); For intracellular FoxP3 staining, cells were collected for fixation and permeabilization, and stained with anti-FoxP3 antibody (Biolegend).

For DNA methylation assay, cells were harvested and washed in PBS, fixed with 4% paraformaldehyde for 15 min at room temperature. After washing in PBS, cells were permeabilized in 0.5% Triton X-100 in PBS for 15 min at room temperature, then washed and treated with 2 N HCl for 30 min at 37°C. Neutralization was performed with 100 mM Tris HCl pH 8.8 for 10 min and followed by extensive washes with 0.05% Tween 20 in PBS. After blocking in 1% BSA and 0.05% Tween 20 in PBS for at least 2 h, cells were incubated with Anti-5-methylcytosine (5-mC) antibody for 1hr, then incubated with 7AAD for 5min for DNA content analysis (Habib et al., 1999).

For histone methylation assay, cells were collected and stained with surface markers, then fixed and permeabilized using the FoxP3 Fixation/Permeabilization Kit (eBioscience). Cells were incubated with a primary monoclonal antibody for 1hr at room temperature, washed, and incubated with the secondary antibody (Alexa-Fluor 647- or Alexa-Fluor 405-conjugated goat anti-mouse Ig antibodies; Invitrogen, France) for 45 min at room temperature. All methylation levels were normalized to total H3 levels. Antibodies were used for detecting the target methylation site (Table 2).Flow cytometry data were acquired on Novocyte (ACEA Biosciences) and were analyzed with FlowJo software (BD Biosciences).

### DNA/RNA/Protein content assay

Cells were collected and stained with surface markers, fixed in 4% paraformaldehyde for 30 min at 4°C, then permeabilized with FoxP3 permeabilization solution (eBioscience). For DNA/RNA content assay, cells were stained with 7AAD for 5 minutes and then stained with pyronin-Y (4ug/ml, PE) for 30 min. Cells were washed and then analyzed by flow cytometer with PerCP channel for 7AAD and PE channel for pyronin-Y. Protein synthesis assay kit (Item No.601100, Cayman) was used for protein content assay. Cells were incubated in O-propargyl-puromycin(OPP) for one hour, fixed, and stained with 5 FAM-Azide staining solutions. Cells were then washed and analyzed by flow cytometer with FITC channel.

### Cell cycle analysis

T cells were incubated with 10 µg/mL BrdU for 1 hour, then surface staining, fixation, and permeabilization according to Phase-Flow Alexa Fluro 647 BrdU Kit (Biolegend). Cells were then stained with antibodies against BrdU to determine the S-phase of the cell cycle, total DNA content was detected by 7AAD labeling. Data were collected on a flow cytometer.

### Western blot analysis

For protein extraction, cells were harvested, lysed, and sonicated at 4 °C in a lysis buffer (50 mM Tris-HCl, pH 7.4, 150 mM NaCl, 0.5% SDS, 5 mM sodium pyrophosphate, protease, and phosphatase inhibitor tablet). Cell lysates were centrifuged at 13,000 × g for 15 min, and the supernatant was recovered. For histone extraction, cells were harvested and resuspended in the TEB buffer, centrifuged, and resuspended the pellet in the extraction buffer. The supernatant was collected (EpiQuik Total Histone Extraction Kit, Epigentek). The protein concentrations were determined using the Pierce™ BCA Protein Assay kit (Thermo Fisher Scientific).

The samples were boiling in NuPAGE® LDS Sample Buffer and Reducing solution (Thermo Fisher Scientific) for 5 min, the proteins/histones were separated by NuPAGE 4-12% Protein Gels (Thermo Fisher Scientific), transferred to PVDF membranes by using the iBlot Gel Transfer Device (Thermo Fisher Scientific), then incubated with primary antibodies (Table 3), followed by incubating with the secondary antibodies conjugated with horseradish peroxidase. Immunoblots were developed on films using the enhanced chemiluminescence technique.

### RNA extraction and RT-qPCR

RNeasy Mini Kit (Qiagen) was used for RNA isolation. Random hexamers and M-MLV Reverse Transcriptase (Invitrogen) was used for cDNA synthesis. BIO-RAD CFX96^TM^ Real-Time PCR Detection System was used for SYBR green-based quantitative PCR. The relative gene expression was determined by the comparative *CT* method, also referred to as the 2^−ΔΔ*CT*^ method. The data were presented as the fold change in gene expression normalized to an internal reference gene (beta2-microglobulin) and relative to the control (the first sample in the group). Fold change = 2^−ΔΔ*C*^_T_ = [(*CT* _gene of interest_- *CT* _internal reference_)] sample A - [(*CT* _gene of interest_- *CT* _internal reference_)] sample B. Samples for each experimental condition were run in triplicated PCR reactions. Primers were used for detecting target genes (Table 4).

### Radioactive tracer-based metabolic activity analysis

The radioactive tracer-based metabolic assay was performed as described previously (Chen et al., 2020). Glycolysis was measured by the generation of ^3^H_2_O from [5-^3^H(N)] D-glucose, fatty acid oxidation was measured by the generation of ^3^H_2_O from [9,10-^3^H] palmitic acid, pentose phosphate pathway was measured by the generation of ^14^CO_2_ from [1-^14^C] D-glucose, glutamine oxidation activity was measured by the generation of ^14^CO_2_ from [U-^14^C]-glutamine. For catabolic activities generation of ^14^CO_2_, five million T cells in 0.5 mL fresh media were dispensed into 7mL glass vials (TS-13028, Thermo) with a PCR tube containing 50μL 0.2N NaOH glued on the sidewall. After adding 0.5 μCi radioactive tracer, the vials were capped using a screw cap with a rubber septum (TS-12713, Thermo) and incubated at 37 °C for 2 hrs. The assay was then stopped by injection of 100μL 5N HCL and kept vials at room temperate overnight to trap the ^14^CO_2_. The NaOH in the PCR tube was then transferred to scintillation vials containing 10mL scintillation solution for counting. A cell-free sample containing 0.5μCi [U-^14^C]-glutamine or [2-^14^C]-pyruvate was included as a background control. For catabolic activities generating ^3^H_2_O, 1μCi radioactive tracer was added to the suspension of one million cells in 0.5mL fresh media in 48 wells, then incubated at 37 °C for 2 hrs. The cells were transferred to a 1.5 mL microcentrifuge tube containing 50μL 5N HCL, placed in 20 mL scintillation vials containing 0.5 mL water with the vials capped and sealed. ^3^H_2_O was separated from other radio-labeled metabolites by evaporation diffusion overnight at room temperature. A cell-free sample containing 1μCi radioactive tracer was included as a background control.

For glutamine flux into the DNA/RNA synthesis, 0.5mM glutamine and 2μCi [U-14C]-glutamine were added to 1.5 X10^7^ cells suspended with glutamine-free medium and cultured for 6hr with IL2. RNA and DNA were isolated with Quick-DNA/RNA™ Miniprep Kit (Cat#: D7005). Radioactivity was measured by liquid scintillation counting and normalized to total DNA or RNA concentration.

### Adoptive cell transfer and *in vivo* proliferation

For homeostatic proliferation assay, isolated naïve CD4^+^ T cells from WT(*Thy1.1^+^*) and *SDHB* cKO(*Thy1.2^+^*) donor mice, cells were mixed with 1:1 ratio and labeled with CFSE. Approximately 1×10^7^ mixed cells suspension in 150 μL PBS were injected via retro-orbital venous sinus into gender-matched *Rag1^-/-^* mice. Five days later, lymph nodes were collected, cell ratio and proliferation were assessed by flow cytometry analysis.

For Ova-antigen driven proliferation assay, isolated naïve CD4^+^ T cells from OT-II-WT (*Thy1.1^+^*) and OT-II-SDHB cKO(*Thy1.2^+^*) donor mice, cells were mixed with 1:1 ratio and labeled with CFSE. Approximately 1×10^7^ mixed cells in 150 μL PBS were injected via retro-orbital venous sinus into gender-matched CD45.1^+^ mice. One day later, host mice were immunized subcutaneously in the hock area of both legs with 50 μL OVA^323-339^ peptides (1 mg/mL, InvivoGen) emulsified with complete Freund’s adjuvant (CFA, InvivoGen). Seven days after immunization, lymph nodes were collected, cell ratio and proliferation were assessed by flow cytometry analysis.

### Experimental Autoimmune Encephalomyelitis (EAE) Induction and Assessment

Mice were immunized with 100 μg of myelin oligodendrocyte glycoprotein (MOG)_35–55_ peptide in CFA (Difco) with 500 μg of Mycobacterium tuberculosis (Difco). Mice were i.p. injected 200 ng of pertussis toxin (PTX, List Biological Laboratories) on the day of immunization and 2 days later. The mice were observed daily for clinical signs and scored as described previously (Shi et al., 2011).

### Histopathology

Mice were euthanized and then were perfused with 25 mL PBS with 2mM EDTA by heart puncture to remove blood from internal organs. Cervical spinal cords were collected and fixed by immersion with 10% neutral buffered formalin solution and decalcified. Tissues were then embedded in paraffin, sectioned, and stained with standard histological methods for H&E staining. Microscopy images were taken using Zeiss Axio Scope A1.

### Stable isotope labeling experiment

#### ^13^C_6_-glucose and ^13^C_5_-glutamine labeling of T cells (Supplemental Figure 1A-C)

CD3^+^ T cells isolated from WT mice were activated for 36 hr, cells were harvested and washed and then seeded with 2*10^6^ cells /mL density in conditional media (RPMI-1640) containing 10 mM ^13^C_6_-glucose or 4 mM ^13^C_5_-Glutamine for 0.5, 2, 6, 12 hours at 37. At the end of each time point, T cells (∼1×10^7^) were washed with PBS 3 times before being snap-frozen.

#### Gas Chromatography-Mass Spectrometry (GC-MS) Sample Preparation and Analysis

GC-MS was performed as previously described(Ratnikov et al., 2015), cell pellets were resuspended in 0.45 ml -20 °C methanol/ water (1:1 v/v) containing 20 µM L-norvaline as internal standard. Further extraction was performed by adding 0.225 ml chloroform, with vortexing and centrifugation at 15,000×g for 5 min at 4 °C. The upper aqueous phase was evaporated under vacuum using a Speedvac centrifugal evaporator. Separate tubes containing varying amounts of standards were evaporated. Dried samples and standards were dissolved in 30 μl 20 mg/ml isobutylhydroxylamine hydrochloride (TCI #I0387) in pyridine and incubated for 20 min at 80°C. An equal volume of N-tertbutyldimethylsilyl-N-methyltrifluoroacetamide (MTBSTFA) (Soltec Ventures) was added and incubated for 60 min at 80 °C. After derivatization, samples and standards were analyzed by GC-MS using an Rxi-5ms column (15 m x 0.25 i.d. x 0.25 μm, Restek) installed in a Shimadzu QP-2010 Plus gas chromatograph-mass spectrometer (GC-MS). The GC-MS was programmed with an injection temperature of 250°C, 1.0 µl injection volume, and a split ratio of 1/10. The GC oven temperature was initially 130°C for 4 min, rising to 250°C at 6°C/min, and to 280°C at 60°C/min with a final hold at this temperature for 2 min. GC flow rate, with helium as the carrier gas, was 50 cm/s. The GC-MS interface temperature was 300°C, and (electron impact) ion source temperature was 200°C, with 70 eV ionization voltage. Fractional labeling from ^13^C substrates and mass isotopomer distributions were calculated as described (Ratnikov et al., 2015). Data from standards were used to construct standard curves in MetaQuant (Bunk et al., 2006), from which metabolite amounts in samples were calculated. Metabolite amounts were corrected for recovery of the internal standard and for ^13^C labeling to yield total (labeled and unlabeled) quantities in nmol per sample and then adjusted per cell number.

#### ^13^C_5_-glutamine labeling of T cells (Figure 2E and Supplemental Figure 5A)

CD3^+^ T Cells (around 1×10^7^ cells/sample) were cultured in conditional media (RPMI-1640) containing 10 mM ^13^C_6_-glucose or 2 mM ^13^C_5_-Glutamine, for 30 hours at 37°C.

#### Capillary electrophoresis-time of flight mass spectrometry (CE-TOFMS) analysis

Metabolome measurements were carried out through a facility service at Human Metabolome Technology Inc., Tsuruoka, Japan. CE-TOFMS measurement was carried out using an Agilent CE Capillary Electrophoresis System equipped with an Agilent 6210 Time of Flight mass spectrometer, Agilent 1100 isocratic HPLC pump, Agilent G1603A CE-MS adapter kit, and Agilent G1607A CE-ESI-MS sprayer kit (Agilent Technologies, Waldbronn, Germany). The systems were controlled by Agilent G2201AA ChemStation software version B.03.01 for CE (Agilent Technologies, Waldbronn, Germany). The metabolites were analyzed using a fused silica capillary (50 μm *i.d.* × 80 cm total length), with commercial electrophoresis buffer (Solution ID: H3301-1001 for cation analysis and H3302-1021 for anion analysis, Human Metabolome Technologies) as the electrolyte. The sample was injected at a pressure of 50 mbar for 10 sec (approximately 10 nL) in cation analysis and 25 sec (approximately 25 nL) in anion analysis. The spectrometer was scanned from *m/z* 50 to 1,000.

#### Capillary electrophoresis–triple quadrupole/time-of-flight mass spectrometry (CE-QqQ/TOFMS) analysis

CD3^+^ T cells isolated from WT and *SDHB* cKO mice were activated for 30hrs. T cells were collected by centrifugation (300 ×*g* at 4°C for 5 min), washed twice with 5% mannitol solution (10 mL first and then 2 mL), then treated with 800 µL of methanol and vortexed for 30 sec to inactivate enzymes.

The cell extract was treated with 550 µL of Milli-Q water containing internal standards (H3304-1002, Human Metabolome Technologies, inc., Tsuruoka, Japan) and vortexed for 30 sec. The extract was obtained and centrifuged at 2,300 ×*g* and 4°C for 5 min, and then 700 µL of the upper aqueous layer was centrifugally filtered through a Millipore 5-kDa cutoff filter at 9,100 ×*g* and 4°C for 180 min to remove proteins. The filtrate was centrifugally concentrated and resuspended in 50 µL of Milli-Q water for CE-MS analysis. Cationic compounds were measured in the positive mode of CE-TOFMS, and anionic compounds were measured in the positive and negative modes of CE-MS/MS. Peaks detected by CE-TOFMS and CE-MS/MS were extracted using an automatic integration software (MasterHands, Keio University, Tsuruoka, Japan and MassHunter Quantitative Analysis B.04.00, Agilent Technologies, Santa Clara, CA, USA, respectively) to obtain peak information, including *m/z*, migration time (MT), and peak area. The peaks were annotated with putative metabolites from the HMT metabolite database based on their MTs in CE and *m*/*z* values determined by TOFMS. The tolerance range for the peak annotation was configured at ±0.5 min for MT and ±10 ppm for *m/z*. In addition, concentrations of metabolites were calculated by normalizing the peak area of each metabolite with respect to the area of the internal standard and by using standard curves, which were obtained by three-point calibrations.

Hierarchical cluster analysis (HCA) and principal component analysis (PCA) were performed by our proprietary software, PeakStat, and SampleStat, respectively. Detected metabolites were plotted on metabolic pathway maps using VANTED (Visualization and Analysis of Networks containing Experimental Data) software.

#### RNA-seq analysis

Total RNA was isolated according to the RNeasy Mini Kit (Qiagen) and treated with DNase I. RNA quality was assessed using the Agilent 2100 Bioanalyzer and RNA Nano Chip kit (Agilent Technologies, CA). RNA-seq libraries were generated using TruSeq Stranded Total RNA with Ribo-Zero Globin Complete kit (Illumina, CA). Ribosomal RNA (rRNA) was removed using target-specific oligos combined with rRNA removal beads. Following rRNA removal, mRNA was fragmented and converted into ds cDNA. The adaptor-ligated cDNA was amplified by limit-cycle PCR. Quality of libraries was determined via Agilent 4200 Tapestation using a High Sensitivity D1000 ScreenTape Assay kit and quantified by KAPA qPCR (KAPA BioSystems). Approximately 60–80 million paired-end 150 bp reads were generated per sample using Illumina HiSeq 4000 platform. For data analysis, each sample was aligned to the GRCm38.p3 assembly of the Mus musculus reference from NCBI using version 2.6.0c of the RNA-seq aligner STAR (Dobin et al., 2013). Features were identified from the GFF file that came with the assembly from Gencode (Release M19). Feature coverage counts were calculated using feature Counts(Liao et al., 2014). Differentially expressed features were calculated using DESeq2 (Love et al., 2014) and significant differential expressions were those with an adjusted *P*-value [i.e., false discovery rate (FDR)] of <0.05.

#### ATAC-seq analysis

ATAC-seq was performed as previously described (Buenrostro et al., 2013) with only minor modifications. 5×10^4^ cells per experiment were first washed with RSB buffer (10 mM Tris-HCl pH 8, 10 mM NaCl, 3 mM MgCl_2_) and gently permeabilized with RSB lysis buffer (10 mM Tris-HCl pH 8, 10 mM NaCl, 3 mM MgCl_2_, 0.1% NP-40) on ice. Cells were suspended in 50 uL of tagmentation master mix prepared from Illumina Tagment DNA TDE1 Enzyme and Buffer Kit components (#20034197), and transposition was performed for 30 minutes at 37°C. Tagmented DNA fragments were isolated using Qiagen MinElute PCR Purification columns prior to library amplification. ATAC-seq libraries were amplified with barcoded Nextera primers for 14 cycles, and excess primers were removed by size selection with AMPure XP beads. Libraries were sequenced on the HiSeq4000 platform running in PEx150bp mode.

The ENCODE ATAC-seq pipeline (https://github.com/ENCODE-DCC/atac-seq-pipeline) with default parameters was used to process ATAC-seq data. First, reads are scanned for adaptor sequences and trimmed with cutadapt (version 2.3). Reads are then mapped to mm10 with bowtie2 (version 2.3.4.3). Properly aligned, non-mitochondrial read pairs were retained for peak calling with MACS2 (version 2.2.4). Differential ATAC peak analysis was completed using DiffBind (Bioconductor) and DESeq2 with an FDR < 0.05. Once differential peaks were called, heatmaps were generated with deeptools (version 3.3.1)(Ramirez et al., 2016), motif analysis was performed using HOMER (version 4.11.1)(Heinz et al., 2010), and nearby genes were identified using GREAT (McLean et al., 2010).

#### CRISPR-Cas 9

All guide RNAs, Alt-R S.p. HIFI Cas9 Nuclease V3, and Alt-R S.p. HIFI electroporation enhancer were purchased from IDT. Mixed 60 pmol of Cas9 protein with 150 pmol of sgRNA and incubated for 20 minutes at room temperature for Cas9/RNP complex formation. Then, activated CD4^+^ T cells were resuspended in P4 Primary Cell buffer (Lonza) and mixed with the RNP complexes and 4uM Electroporation Enhancer in P4 Primary Cell buffer (Lonza), transferred the cell/CRISPR mixture to the bottom hole of the wells of the Lonza nucleofector strip for electroporation by using Lonza nucleofector (Program CM137), electroporated cells were recovered in T cell medium for 2 hours prior to activation with anti-CD3/CD28 antibodies, or under T_H_1 or T_H_17 polarization condition. Duplexes of two separate guides per target gene were used: Mm.Cas9.PRDM1.1AA (ACACGCTTTGGACCCCTCAT), Mm.Cas9.PRDM1.1AB (CATAGTGAACGACCACCCCT).

### Statistical analysis

Statistical analysis was conducted using the GraphPad Prism software (GraphPad Software, Inc.). *P* values were calculated with two-way ANOVA for the EAE experiments. Unpaired two-tail Student’s t-test, multiple comparisons of one/ two-way ANOVA, or Dunnett LSD test was used to assess differences in other experiments. *P* values smaller than 0.05 were considered significant, with *p* < 0.05, *p* < 0.01, *p* < 0.001, and *p* < 0.0001 indicated as *, **, ***, and ****, respectively.

## Figure Legends

Supplemental figure legends are given in Supplementary Information.

