## Supplementary material for "Succinate dehydrogenase/complex II is critical for metabolic and epigenetic regulation of T cell proliferation and inflammation": S figure

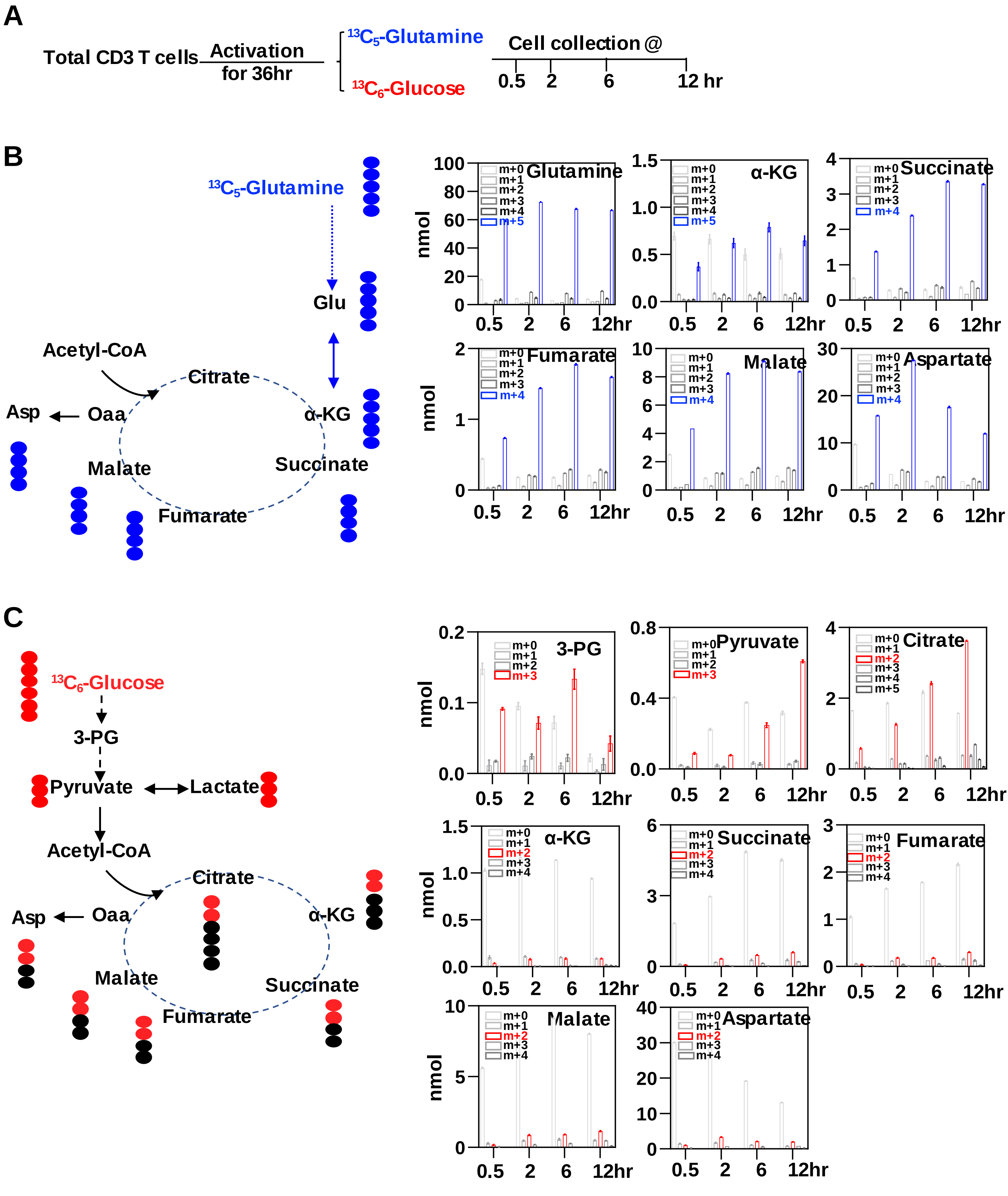


**Supplemental figure 1. The TCA cycle allocates carbon input from glucose and glutamine to support T cell growth and proliferation.**

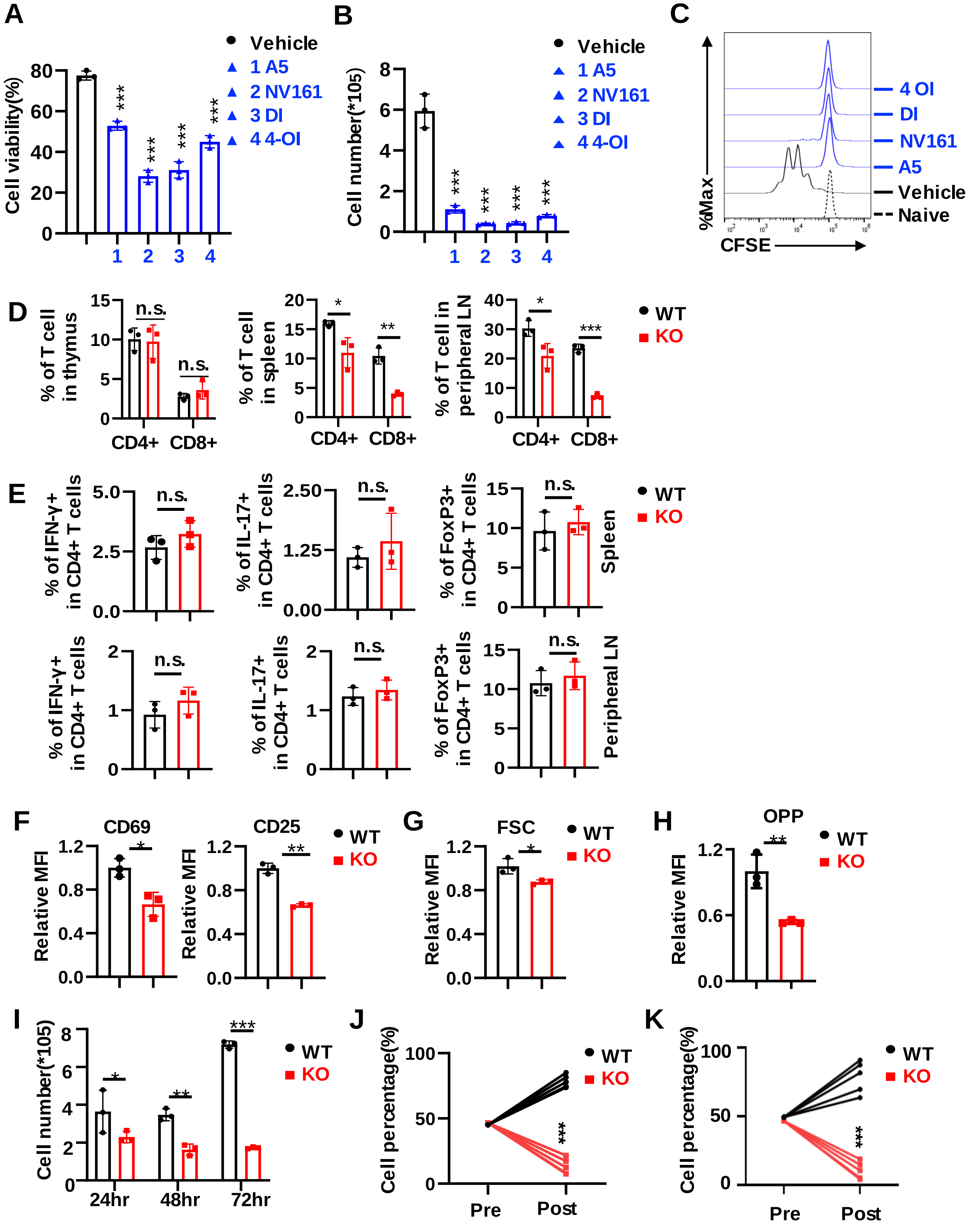


**Supplemental figure 2. Complex II/SDHB is required for T cell activation and development.**

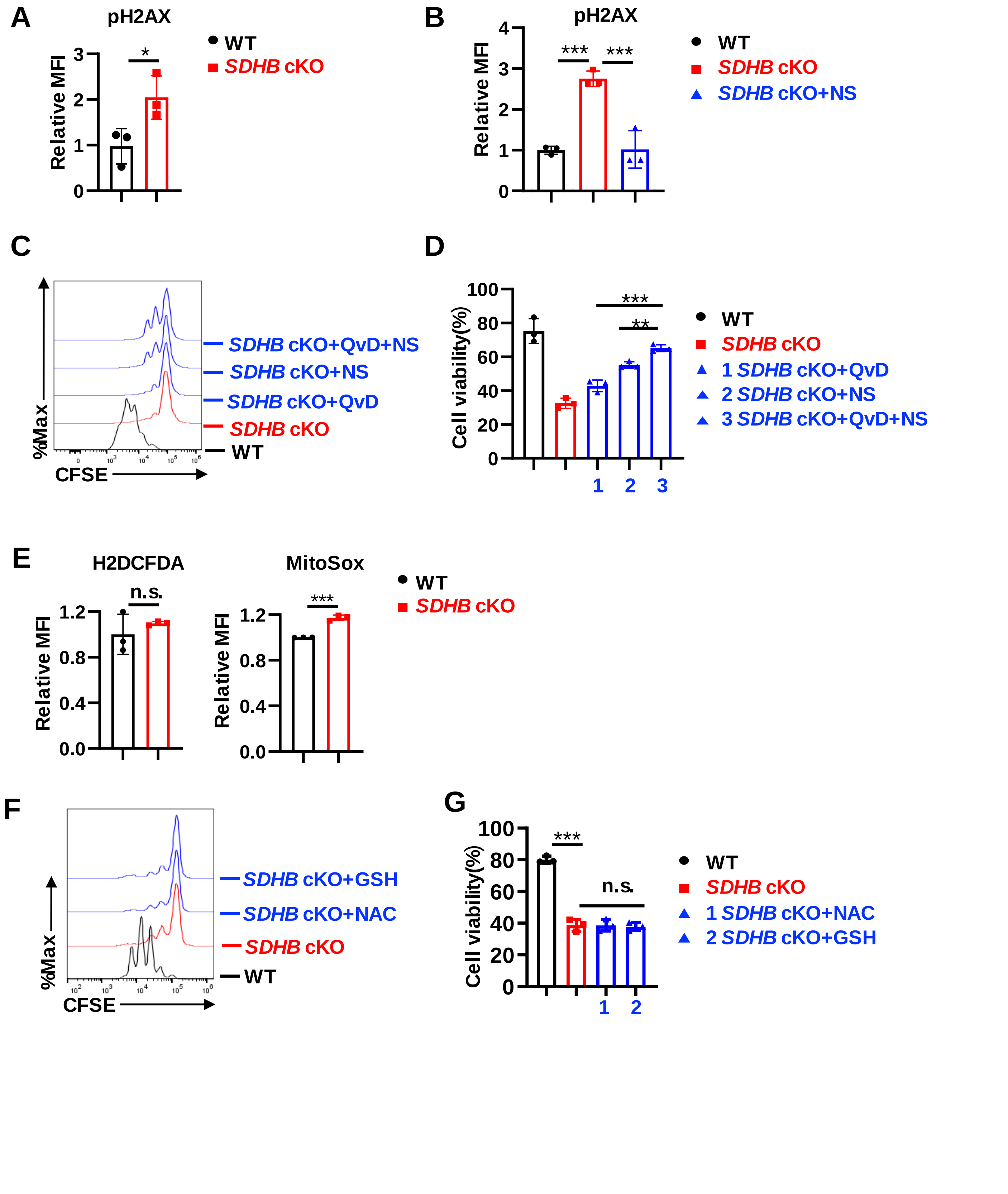


**Supplemental figure 3. SDHB deficiency induces DNA damage and moderately increases ROS production.**

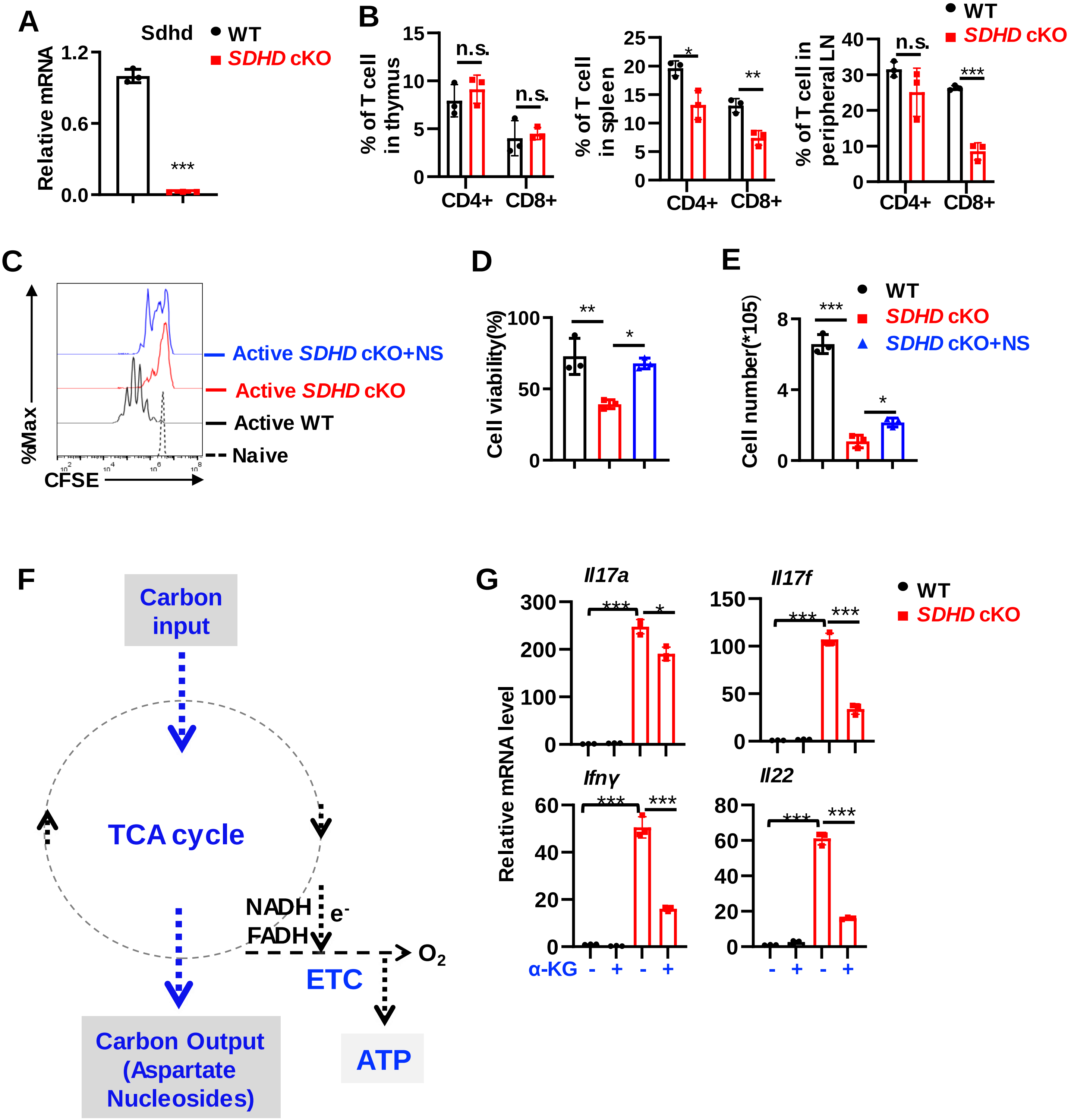


**Supplemental figure 4. SDHD deficiency phenocopies SDHB deficiency in T cell.**

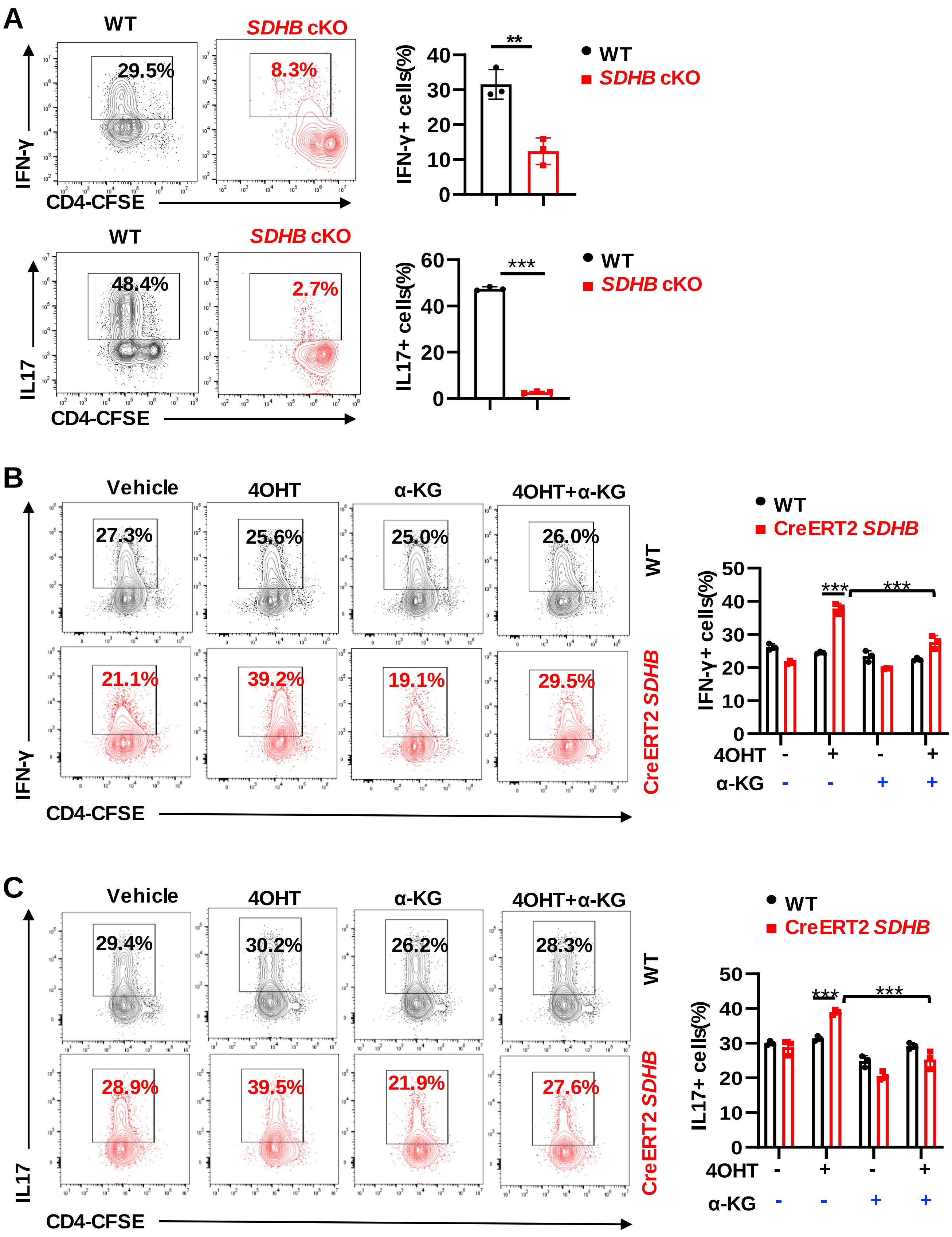


**Supplemental figure 5. Acute deletion of SDHB enhances $T_H1$ and $T_H17$ differentiation.**

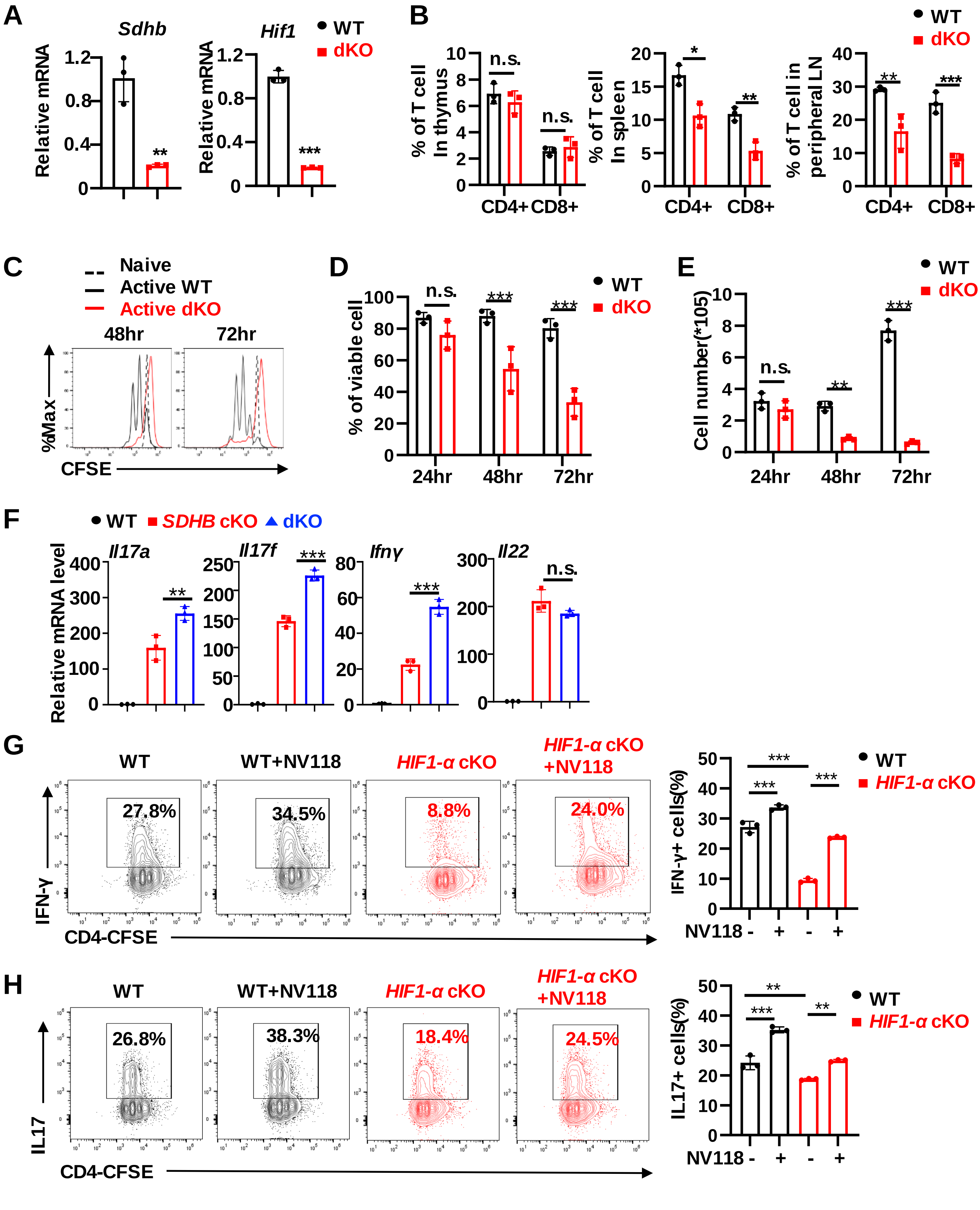


**Supplemental figure 6. HIF1α is dispensable for pro-inflammatory gene signature caused by SDHB deficiency.**

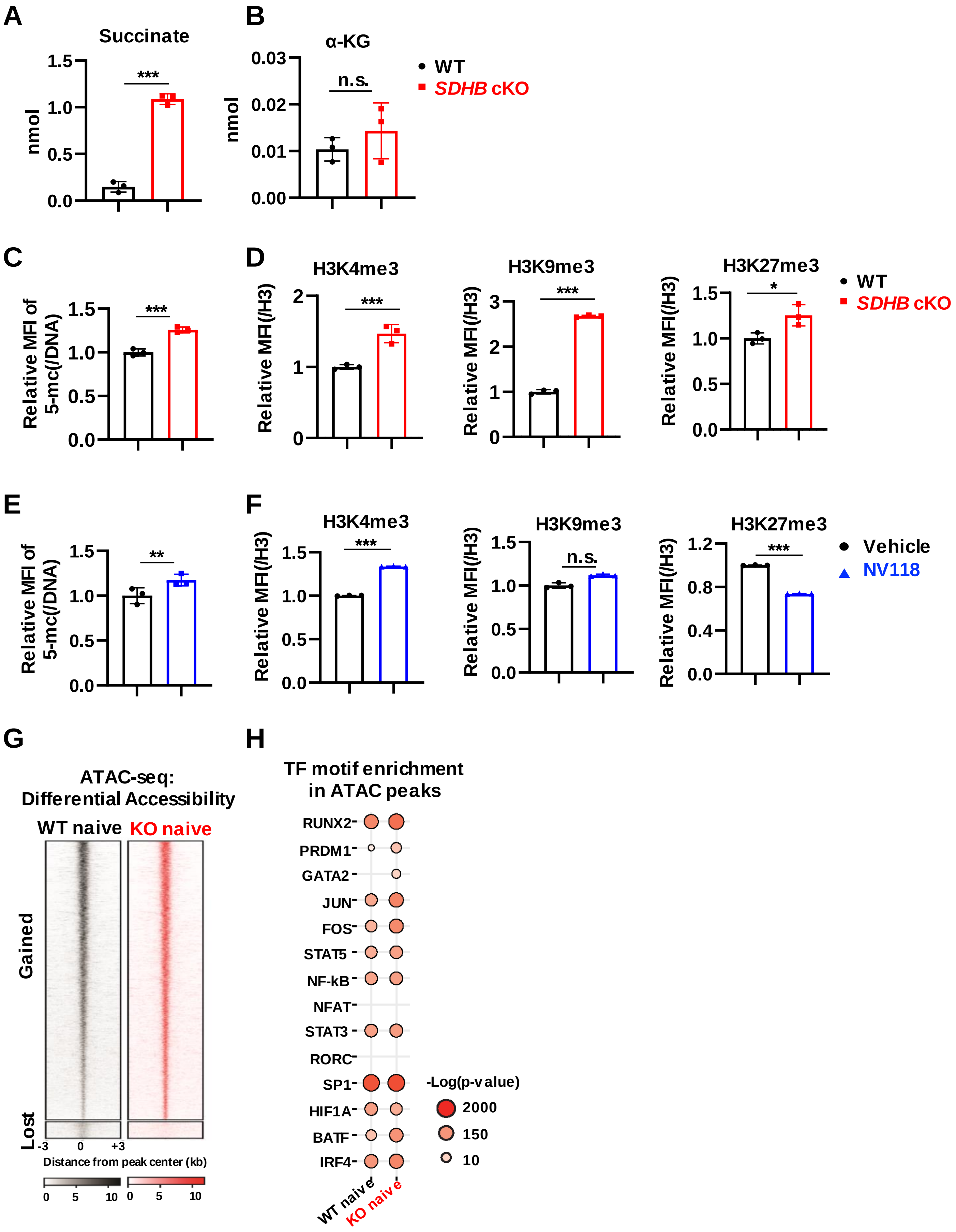


**Supplemental figure 7. Succinate accumulation impacts T cell epigenome.**
