## Supplementary material for "Succinate dehydrogenase/complex II is critical for metabolic and epigenetic regulation of T cell proliferation and inflammation": SI

**Supplemental figure legends**

**Supplemental figure 1. The TCA cycle allocates carbon input from glucose and glutamine to support T cell growth and proliferation.**

**(A)** The experimental procedure of isotope ^13^C_5_-Glutamine and ^13^C_6_-Glucose labeling and mass spectrometry approach. WT total T cells were activated for 36hr, then incubated in ^13^C_5_-Glutamine or ^13^C_6_-Glucose medium, cells were collected at the indicated time points, metabolites were extracted and analyzed using GC-MS. **(B)** Diagram of putative catabolic routes of ^13^C_5_-Glutamine in T cells (left panel), metabolites were present in the right panel, numbers in the X-axis represent indicated time points, bar with different colors represent those of ^13^C atoms in given metabolites, numbers in the Y-axis represent the levels of the metabolites (nmol). Data represent mean±SEM of (n=3) for each group. **(C)** Diagram of putative catabolic routes of ^13^C_6_-Glucose in T cells (left panel), metabolites were present in the right panel, numbers in the X-axis represent indicated time points, bar with different colors represent those of ^13^C atoms in given metabolites, numbers in the Y-axis represent the levels of the metabolites (nmol). Data represent mean±SEM of (n=3) for each group.

**Supplemental figure 2. Complex II/SDHB is required for T cell activation and development.**

**(A-C)** WT CD4^+^ T cells were activated and exposed to SDH/complex II inhibitors including atpenin A5 (A5, 100nM), diacetoxymethyl malonate (NV161,100μM), dimethyl-itaconate (DI, 0.5mM), or 4-octyl-itaconate (4OI, 500μM).  **(A)** The statistical analysis of cell viability of activated CD4^+^ T cells at 72hr was assessed by 7AAD staining. **(B)** Cell number was calculated by cell counter. Data represent of 3 independent experiments, ****p* < 0.001, one-way ANOVA. **(C)** Cell proliferation of naïve and activated CD4^+^ T cells at 72hr was determined by CFSE dilution. Data represent at least 3 independent experiments. **(D-E)** Distribution of indicated T cell subsets in the thymus, spleen, and peripheral lymph node (LN) were determined by cell surface and intracellular markers by flow cytometry (mean±SEM, n=3). Data represent of 3 independent experiments, **p* < 0.05, ***p* < 0.01, ****p* < 0.001, one-way ANOVA. **(F)** CD4^+^ T cell activation at 24hr was determined by the cell surface expression of CD25(left panel) and CD69(right panel), mean fluorescence intensity (MFI) was analyzed, ****p* < 0.001, student’s *t* test. **(G)** CD4^+^ T cell size at 24hr was measured by the forward scatter (FSC), MFI was analyzed **p* < 0.05, student’s *t* test. **(H)** CD4^+^ T cell protein content at 24hr was determined by the OPP staining, MFI was analyzed, **p* < 0.05, student’s *t* test. **(I)** Cell number of activated CD4^+^ T cells at 24hr, 48hr and 72hr with indicated genotypes was measured by the cell counter, **p* < 0.05, ***p* < 0.01, ****p* < 0.001, one-way ANOVA. **(J)** CD4 T cell ratios of *in vivo* competitive proliferation before and after adoptive transfer were evaluated by surface staining of isogenic markers. **(K)** CD4 T cell ratios of *in vivo* antigen-specific competitive proliferation before and after adoptive transfer were evaluated by surface staining of isogenic markers (mean±SEM, n=5). Data represent 3 independent experiments. ****p* < 0.001, two-way ANOVA

**Supplemental figure 3. SDHB deficiency induces DNA damage and moderately increases ROS production.**

**(A-B)** CD4^+^T cells from indicated genotypes without **(A)** or with mixture NS **(B)** were activated for 24 hr, DNA damage was analyzed by flow cytometry with H2A.X Phospho (Ser139) Antibody, the relative MFI of γH2AX was determined, MFI was analyzed, **p* < 0.05, ****p* < 0.001. **(C-D)** *SDHB* cKO CD4^+^ T cells were supplemental of mixture NS, QVD-OPh, NS+ QVD-OPh, cell proliferation of activated CD4^+^ T cells at 72hr was determined by CFSE dilution **(C)**, cell viability at 72 hr was calculated based on the 7AAD staining **(D)**, ***p* < 0.01, ****p* < 0.001, one-way ANOVA. (**E**) CD4^+^ T cells from indicated genotypes were activated for 24 hr, intracellular ROS and mitochondrial ROS were measured by the H2DCFDA staining or MitoSOX™ Red staining, MFI was analyzed, ****p* < 0.001, student’s *t* test. **(F-G)** CD4^+^ T cells from indicated genotypes were incubated with or without 1mM NAC or 1mM GSH EE. Cell proliferation of activated CD4^+^ T cells at 72hr was determined by CFSE dilution **(F)**, cell viability at 72 hr was calculated based on the 7AAD staining **(G)**. Data represent of 3 independent experiments, ****p* < 0.001, one-way ANOVA.

**Supplemental figure 4. SDHD deficiency phenocopies SDHB deficiency in T cell.**

**(A)** SDHD mRNA levels in activated T cells was determined by qPCR (mean±SEM, n=3), ****p* < 0.001, student’s *t* test. **(B)** Distribution of indicated T cell subsets in the thymus, spleen, and peripheral lymph node (LN) were determined by cell surface and intracellular markers by flow cytometry (mean±SEM, n=3). Data represent of 3 independent experiments, **p* < 0.05, ***p* < 0.01, ****p* < 0.001, one-way ANOVA. **(C-E)** *SDHD* cKO CD4^+^ T cells were supplemental of a mixture of nucleosides (NS), cell proliferation of activated CD4^+^ T cells at 72hr was determined by CFSE dilution **(C)**, cell viability at 72 hr was calculated based on the 7AAD staining **(D)**, cell number at 72 hr was measured by a cell counter **(E).** Data represent of 3 independent experiments, **p* < 0.05, ***p* < 0.01, ****p* < 0.001, one-way ANOVA. **(F)** Schematic view of carbon input and output through the TCA cycle in SDH dysfunction models. **(G)** CD4 T cells were activated for 36hr with or without α-KG, **mRNA levels of indicated genes were measured by qPCR** (mean±SEM, n=3)**,** **p* < 0.05, ****p* < 0.001, one-way ANOVA.

**Supplemental Figure 5. Acute deletion of SDHB enhances T_H_1 and T_H_17 differentiation.**

**(A)** CD4^+^T cells from WT and *SDHB* cKO mice were polarized toward T_H_1 and T_H_17 lineages for 72 hours. The indicated proteins were quantified by intracellular staining, cell proliferation was determined by CFSE staining. Data represent three independent experiments, ****p*<0.001, student’s *t* test. **(B-C)** CD4^+^T cells from WT and tamoxifen-induced Cre recombinase (CreERT2) *SDHB* cKO mice were polarized toward T_H_1 **(B)** and T_H_17 **(C)** lineages with indicated treatment for 72 hours. The indicated proteins were quantified by intracellular staining, cell proliferation was determined by CFSE staining. Data represent three independent experiments, ****p*<0.001, one-way ANOVA.

**Supplemental Figure 6. HIF1𝜶 is dispensable for pro-inflammatory gene signature caused by SDHB deficiency.**

**(A)** mRNA levels of indicated genes were measured by the qPCR (mean±SEM, n=3), ***p*<0.01, ****p*<0.001, student’s *t* test. **(B)** Distribution of indicated T cell subsets in the thymus, spleen, and peripheral lymph node (LN) were determined by cell surface markers by flow cytometry (mean±SEM, n=3). Data represent of 3 independent experiments, **p* < 0.05, ***p* < 0.01, ****p* < 0.001, one-way ANOVA. **(C-E)** Cell proliferation of naïve and activated CD4^+^ T cells at 72hr was determined by CFSE dilution **(C)**, cell viability of activated CD4^+^ T cells at 72hr was assessed by 7AAD staining, the statistical analysis of cell viability (mean ± SEM) **(D)**, cell number was calculated by a cell counter **(E)**. Data represent of 3 independent experiments, ***p* < 0.01, ****p* < 0.001, one-way ANOVA. **(F)** CD4 T cells from indicated genotypes were activated for 36hr, mRNA levels of indicated genes were determined by qPCR (mean±SEM, n=3), ***p* < 0.01, ****p* < 0.001, one-way ANOVA. **(G-H)** CD4^+^T cells from WT and *HIF1-𝜶* cKO mice were polarized toward T_H_1 **(G)** and T_H_17 **(H)** lineages with indicated treatment for 72 hours. The indicated proteins were quantified by intracellular staining, cell proliferation was determined by CFSE staining. Data represent three independent experiments, ***p* < 0.01, ****p* < 0.001, one-way ANOVA.

**Supplemental figure 7. Succinate accumulation impacts T cell epigenome.**

**(A-B)** Indicated metabolites in naive T cells from indicated genotypes were determined, ****p*<0.001, student’s *t* test. **(C)** DNA methylation levels in naive T cells from indicated genotypes were determined by 5-mc staining through flow cytometry. **(D)** Indicated histone methylation levels in naive T cells from indicated genotypes were determined by intracellular staining through flow cytometry, **p*<0.05, ****p*<0.001, student’s *t* test. **(E)** WT CD4^+^ T cells were incubated with NV118 for 72hr in naïve condition, DNA methylation levels were determined by 5-mc staining through flow cytometry, **(F)** indicated histone methylation levels in naive T cells with indicated treatment were determined by intracellular staining through flow cytometry, *******p*<0.01, ****p*<0.001, student’s *t* test. (**G**) Differential chromatin accessibility was measured by ATAC-seq in naïve CD4 T cells between indicated genotypes (n = 3 replicates for each genotype), identifying 5,086 sites with accessibility gain and 346 sites with accessibility loss in *SDHB* cKO T cells. (**H**) Motif analysis was performed in differential ATAC-seq peaks (WT vs. *SDHB* cKO ) in naïve CD4 T cells. Motif enrichment for transcription factors involved in T cell activation and inflammation are provided.
