## Supplementary material for "Succinate dehydrogenase/complex II is critical for metabolic and epigenetic regulation of T cell proliferation and inflammation": table1

| **Name** | **Cat#** | **Vendor** |
| --- | --- | --- |
| InVivoMAb anti-mouse CD3 | BE0001-1 | BioXcell |
| InVivoMAb anti-mouse CD28 | BE0015-1 | BioXcell |
| InVivoMAb anti-mouse IFNγ | BP0055 | BioXcell |
| InVivoMAb anti-mouse IL-4 | 11B11 | BioXcell |
| InVivoMAb anti-mouse IL-2 | BE0043-1 | BioXcell |
| Recombinant Murine IL-12 p70 | 210-12 | Peprotech |
| InVivoMAb anti-mouse IL-2 | 212-12 H1111 | Peprotech |
| Recombinant Murine IL-7 | 217-17 | Peprotech |
| Recombinant Murine IL-6 | 216-16 | Peprotech |
| Adenosine | AC164040050 | Thermo Fish Scientific |
| Uridine | U3003 | Sigma-Aldrich |
| Inosine | I4125 | Sigma-Aldrich |
| Cytidine | C4654 | Sigma-Aldrich |
| Guanosine | G6264 | Sigma-Aldrich |
| Thymidine | T1895 | Sigma-Aldrich |
| Q-VD-OPH(QvD) | 1135695-98-5 | BOC science |
| Atpenin A5 | 119509-24-9 | Cayman |
| polyketides NV161 | Cpd ID 01-161-S2; Vial ID BION10476 | Isomerase therapeutics Ltd. Cambridge |
| polyketides NV118 | Cpd ID 01-118-s3; Vial ID BION10474 | Isomerase therapeutics Ltd. Cambridge |
| Dimethyl itaconate | 592498 | Sigma-Aldrich |
| 4-octyl-itaconate | 3133-16-2 | Sigma-Aldrich |
| Dimethyl 2-oxogluterate(Dim α-KG) | 349631 | Sigma-Aldrich |
| (Z)-4-Hydroxytamoxifen(4OHT) | 68047-06-3 | Sigma-Aldrich |
| GDH1 Inhibitor, R162 - Calbiochem | 5380980001 | Sigma-Aldrich |
| NAC | A7250 | Sigma-Aldrich |
| Glutathione reduced ethyl este(GSH EE) | G1404 | Sigma-Aldrich |

**Table 1. Cell culture related antibodies, cytokines, inhibitors, chemicals**
