## Supplementary material for "Succinate dehydrogenase/complex II is critical for metabolic and epigenetic regulation of T cell proliferation and inflammation": table2

| **Name** | **Cat#** | **Vendor** |
| --- | --- | --- |
| Pacific Blue™ anti-mouse CD4 Antibody | 100428 | BioLegend |
| APC anti-mouse CD4 Antibody | 100516 | BioLegend |
| PE/Cyanine7 anti-mouse CD4 Antibody | 100422 | BioLegend |
| APC/Cyanine7 anti-mouse CD8a Antibody | 100714 | BioLegend |
| APC anti-mouse TCR β chain Antibody | 109211 | BioLegend |
| PE/Cyanine7 anti-mouse CD90.1 (Thy-1.1) Antibody | 202518 | BioLegend |
| APC/Cyanine7 anti-mouse CD90.2 Antibody | 105328 | BioLegend |
| APC anti-mouse CD90.2 (Thy-1.2) Antibody | 140311 | BioLegend |
| PE/Cyanine7 anti-mouse CD45.1 Antibody | 110730 | BioLegend |
| APC/Cyanine7 anti-mouse CD45.2 Antibody | 109824 | BioLegend |
| FITC anti-mouse CD3 Antibody | 100204 | BioLegend |
| PE-Cy™7 Hamster Anti-Mouse CD69 | 552879 | BD Pharmingen™ |
| CD25 Monoclonal Antibody (PC61.5), PE | 12-0251 | eBioscience™ |
| APC anti-mouse IFN-γ Antibody | 505810 | BioLegend |
| PE/Cyanine7 anti-mouse IFN-γ Antibody | 505826 | BioLegend |
| APC anti-mouse IL-17A Antibody | 506916 | BioLegend |
| PE/Cyanine7 anti-mouse IL-17A Antibody | 506922 | BioLegend |
| Alexa Fluor® 488 Donkey anti-rabbit IgG Antibody | 406416 | BioLegend |
| Alexa Fluor® 647 Donkey anti-rabbit IgG Antibody | 406414 | BioLegend |
| PE/Cyanine7 anti-H2A.X Phospho (Ser139) Antibody | 613419 | BioLegend |
| Alexa Fluor® 647 anti-mouse FOXP3 Antibody | 126407 | BioLegend |
| Anti-5-methylcytosine (5-mC) Antibody (APC) | AC16-0073-03 | Abcore |
| Blimp-1 Monoclonal Antibody (5E7), Alexa Fluor 488 | 53-9850-82 | eBioscience™ |
| [Tri-Methyl-Histone H3 (Lys4) (C42D8) Rabbit mAb](https://www.cellsignal.com/products/primary-antibodies/9751) | #9751 | Cell Signaling |
| Tri-Methyl-Histone H3 (Lys9) (D4W1U) Rabbit mAb | #13969 | Cell Signaling |
| [Tri-Methyl-Histone H3 (Lys27) (C36B11) Rabbit mAb](https://www.cellsignal.com/products/primary-antibodies/tri-methyl-histone-h3-lys27-c36b11-rabbit-mab/9733) | #9733 | Cell Signaling |
| [Histone H3 (D1H2) XP® Rabbit mAb](https://www.cellsignal.com/products/primary-antibodies/histone-h3-d1h2-xp-rabbit-mab/4499) | #4499 | Cell Signaling |
| Pyronin Y | 92-32-0 | Sigma-Aldrich |
| APC anti-BrdU Antibody | 364114 | BioLegend |
| MitoSOX™ Red Mitochondrial Superoxide Indicator | M36008 | Invitrogen™ |
| Carboxy-H2DFFDA | C13293 | Invitrogen™ |
| 7-AAD Viability Staining Solution | 420404 | BioLegend |

**Table 2. Cell staining antibodies and dyes**
