## Supplementary material for "Succinate dehydrogenase/complex II is critical for metabolic and epigenetic regulation of T cell proliferation and inflammation": table4

| **Gene** | **primer sequences forward** | **primer sequences reverse** |
| --- | --- | --- |
| Sdhb | ATTTACCGATGGGACCCAGAC | GTCCGCACTTATTCAGATCCAC |
| Sdhd | TGGTCAGACCCGCTTATGTG | GGTCCAGTGGAGAGATGCAG |
| Ifng | ATGAACGCTACACACTGCATC | CCATCCTTTTGCCAGTTCCTC |
| Il17a | TTTAACTCCCTTGGCGCAAAA | CTTTCCCTCCGCATTGACAC |
| Il17f | TGCTACTGTTGATGTTGGGAC | AATGCCCTGGTTTTGGTTGAA |
| Il22 | ATGAGTTTTTCCCTTATGGGGAC | GCTGGAAGTTGGACACCTCAA |
| Hif1a | AGCTTCTGTTATGAGGCTCACC | TGACTTGATGTTCATCGTCCTC |
| Tubulin | TTCTGGTGCTTGTCTCACTGA | CAGTATGTTCGGCTTCCCATTC |
| Gata2 | CACCCCGCCGTATTGAATG | CCTGCGAGTCGAGATGGTTG |
| Runx2 | AACGATCTGAGATTTGTGGGC | CCTGCGTGGGATTTCTTGGTT |
| Tbx6 | ATGTACCATCCACGAGAGTTGT | GGTAGCGGTAACCCTCTGTC |
| Zeb2 | ATTGCACATCAGACTTTGAGGAA | ATAATGGCCGTGTCGCTTCG |
| Kif4 | AGGTGAAGGGGATTCCCGTAA | AAACACGCCTTTTATGAGTGGA |
| Spib | AGGAGTCTTCTACGACCTGGA | GAAGGCTTCATAGGGAGCGAT |
| Ebf1 | GCATCCAACGGAGTGGAAG | GATTTCCGCAGGTTAGAAGGC |
| Mef2c | GTCAGTTGGGAGCTTGCACTA | CGGTCTCTAGGAGGAGAAACA |
| Bhlhe41 | TGTGTAAACCCAAAAGGAGCTT | TGTTCGGGCAGTAAATCTTTCAG |
| Rorc | CGCGGAGCAGACACACTTA | CCCTGGACCTCTGTTTTGGC |
| Prdm1 | TTCTCTTGGAAAAACGTGTGGG | GGAGCCGGAGCTAGACTTG |
| Irf4 | TCCGACAGTGGTTGATCGAC | CCTCACGATTGTAGTCCTGCTT |
| Stat3 | CAATACCATTGACCTGCCGAT | GAGCGACTCAAACTGCCCT |
| Batf | CTGGCAAACAGGACTCATCTG | GGGTGTCGGCTTTCTGTGTC |
| Bach1 | TGGTGAGAGTGCGGTATTTGC | GTCAGTCTGGCCTACGATTCT |

**Table 4. RT-qPCR primers**
